# Protein-Solvent Shape Complementarity as a Unifying Principle in Excipient-Mediated Protein Thermal Stability

**DOI:** 10.64898/2026.06.12.731979

**Authors:** Jonathan W. P. Zajac, Praveen Muralikrishnan, Xianci Zeng, Caryn L. Heldt, Sarah L. Perry, Sapna Sarupria

## Abstract

Excipient effects on protein stability are critical for biological formulations, yet their selection remains largely empirical. Here, we use molecular dynamics simulations to define unifying metrics of protein-excipient interactions at atomistic resolution. Enhanced sampling simulations of fast-folding miniproteins, including Trpzip, WAAAH-helix (an alanine-rich *α*-helix), and Trp-Cage, were performed to capture folding transitions across diverse excipient conditions. We identified a general stabilization mechanism based on shape complementarity between protein networks and surrounding solvent networks. Stabilizing excipients were found to form solvent structures that preferentially complement each protein, as well as residues central to known folding pathways. This framework enables a unifying approach to mechanism-based excipient selection across diverse protein and solvent chemistries. More broadly, by treating protein and solvent as dynamically coupled partners, it provides a transferable strategy for understanding solvent-mediated effects in complex molecular systems.

---

Developing biologics with long shelf lives poses a grand challenge in formulation design. The primary constituent of many biologics, such as virus-like particles (VLPs), vaccines, gene therapy products, or monoclonal antibodies, is proteins. Proteins are susceptible to degradation in formulations due to both chemical and physical denaturation. ^1^ The temperature dependence of proteins is a key source of physical denaturation in formulations, as proteins unfold at both very low and high temperatures.^2,3^ Due to this temperature sensitivity, most biologics are maintained within a supply chain known as a “cold chain”.^4,5^ The cold chain stores biologics at refrigerated temperatures, though disruptions can cause potency loss of the product.^6^ Strategies that obviate the need for the cold chain are thus an attractive avenue for designing stable biological formulations.

Inspired by intracellular osmolyte production, excipient innovation is a promising strategy in the development of formulations independent of cold storage. This process involves the addition of small molecules to formulations to improve stability.^7^ The excipient selection step of the formulation design process is generally high-throughput and empirical due to the large number of possible excipients and frequent usage of multiple excipients in a single formulation.^8^ The trial-and-error nature of this process bottlenecks formulation development, expensive in terms of both money and time.^9^

A paradigm shift is currently underway in the design of stable biologics, from trial-and-error screening to rational design based on mechanistic understanding of protein-excipient relationships. ^10–13^ Recent developments of resource-saving design strategies have illustrated that systematic and effective formulation development does not depend on a single physical quantity.^14–16^ Instead, effective design strategies balance multiple properties, such as conformational stability (folding/unfolding) and colloidal stability (aggregation).^17^ Recent work towards understanding protein-excipient mechanisms points to the local protein-solvent environment as an integral factor in determining an excipient’s stabilizing propensity.^18–34^ Properties including the local accumulation or depletion of excipients, the relative hydration state of the protein, hydration shell dynamics, and hydrogen bonding networks have all been implicated in native state protein stability. Importantly, each of these properties is dictated by a fine balance of excipient-water-protein interactions and therefore depends on both the identity of the excipient and the protein.^35–38^ Due to this complexity, there is an unmet need for the development of a unifying framework that is transferable across diverse excipient-protein combinations while accurately describing biomolecular stability.

To address this challenge and develop unified mechanistic descriptors capturing excipient effects on protein stability, we studied stability of three miniproteins across 11 excipient solutions (Fig. S1). The selected excipients cover several distinct features, including positive charge (arginine, ARG; lysine, LYS), negative charge (glutamate, GLU), polar side chain (serine, SER), aromatic side chain (phenylalanine, PHE), hydrophobic side chain (leucine, LEU), homodisaccharide (trehalose, THL), heterodisaccharide (sucrose, SCRS), monosaccharide (*α*-glucose, AGLC), and sugar alcohol (sorbitol, SOR; glycerol, GOL).

The three miniproteins were selected for their fast-folding times, experimentally known melting temperatures (Table 1), and distinct structural features. Trpzip is stabilized through Trp-stacking interactions and zipper-like Thr-Thr hydrogen bonds, serving as a model for *β*-hairpin formation.^39^ WAAAH-Helix forms an *α*-helix through its Ala-rich (AAARA)_3_ motif and has been used as a model to understand helix-forming folding pathways. ^40^ Trp-Cage is a well-characterized model for protein folding,^41–44^ consisting of an N-terminal helix that forms either before or alongside the collapse of a hydrophobic-lined cage surrounding a central Trp6 residue. An Arg9-Asp16 salt bridge forms after the folding transition state, which further stabilizes the native state.

**Table 1:**
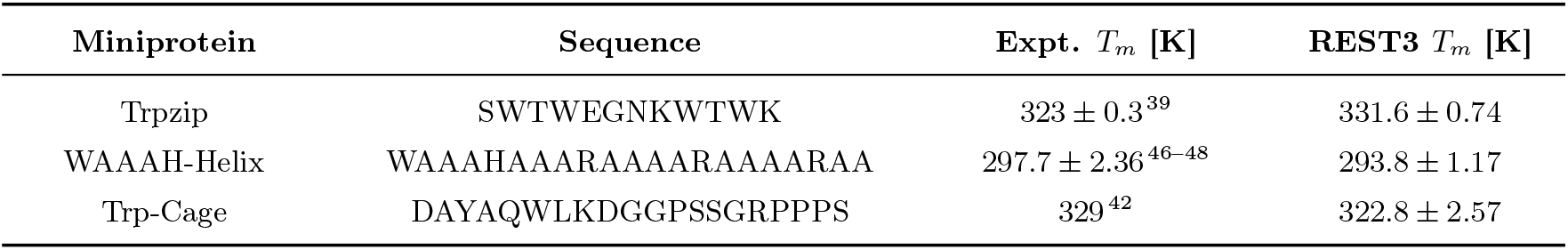
Miniprotein sequences, experimentally (expt.) determined melting temperatures, and melting temperatures obtained from REST3 simulations in the present work.

| Miniprotein | Sequence | Expt. $T_m$ [K] | REST3 $T_m$ [K] |
| --- | --- | --- | --- |
| Trpzip | SWTWEGNKWTWK | $323 \pm 0.3$<sup>39</sup> | $331.6 \pm 0.74$ |
| WAAAH-Helix | WAAAHAAARAAAARAAAARAA | $297.7 \pm 2.36$<sup>46-48</sup> | $293.8 \pm 1.17$ |
| Trp-Cage | DAYAQWLKDGGPSSGRPPPS | $329$<sup>42</sup> | $322.8 \pm 2.57$ |

We performed Replica Exchange with Solute Tempering 3 (REST3) to efficiently explore the conformational landscapes of miniprotein folding/unfolding. ^45^ REST3 uses Hamiltonian rescaling to simulate different regions of the system under different effective temperatures. REST3 improves sampling efficiency by treating solute-solvent van der Waals (vdW) interactions as an independent, tunable parameter scaled according to *κ*_*m*_. For Trp-Cage and Trpzip, we applied *κ*_*m*_ = 1 at lower temperature replicas (indices *m* ≤ 3) and *κ*_*m*_ = 1 + 0.005 ^*^ (*m* − 3) for higher temperature replicas (indices *m* ≥ 4). To recover the experimental *T*_*m*_ of WAAAH-Helix, we found it was necessary to weaken protein-solvent interactions at lower temperatures (Fig. S2). This was achieved by setting *κ*_*m*_ according to *κ*_*m*_ = 1 + 0.025(*m* − 12).

To monitor the folding progress of the miniproteins, we used the fraction of native contacts,^49^ Q. Q was computed for all miniproteins as 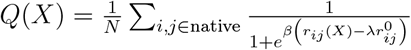, where *N* is the total number of native contacts, *r*_*ij*_(*X*) is the distance between heavy atoms *I* and *j* in configuration *X*, 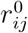 is the distance between heavy atoms *i* and *j* in the native state, *β* is a smoothing parameter taken to be 0.5 nm^*−*1^, and *λ* is the contact tolerance relative to native distance, taken to be 1.8. Native contacts and their distances were assigned according to the energy minimized crystal structures of Trpzip (Trpzip1 variant; PDB: 1LE0)^39^ and TrpCage (tc10b variant; PDB: 2JOF).^42^ As there is no experimentally available crystal structure of WAAAH-Helix, we considered only the contacts formed among residues central to the *α*-helix (residues 6–16) in the calculation of *Q* for this protein. For each temperature replica, the ensemble-averaged fraction of native contacts was computed as 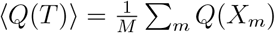, where *M* is the number of configurations sampled at temperature *T*. At each effective temperature *T*, the folded fraction was approximated as *f* (*T*) ≈ ⟨*Q*(*T*)⟩. The free energy of protein folding, Δ*G*(*T*) was calculated as 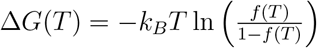. Δ*G* was fit to the analytical expression for a two-state folding process, 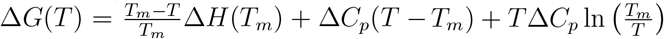, to estimate the melting temperature, *T*_*m*_. The melting temperatures calculated from the simulations are consistent with the reported experimental values (Table 1).

An effective temperature range from *T*_0_ to *T*_*f*_ encompassing the experimentally known melting temperatures was used for all REST3 simulations. For Trpzip and Trp-Cage, *T*_0_ = 300 K and *T*_*f*_ = 510 K, and for WAAAH-Helix, *T*_0_ = 260 K to *T*_*f*_ = 470 K. Effective temperatures were distributed according to a quadratic function such that higher effective temperatures were spaced further apart. Simulations were run until the probability distribution of the fraction of native contacts, *P* (*Q*), was similar in two evenly sized blocks of simulation time (Figs. S3–S5). REST3 replicas were performed for 200-400 ns, Trpzip; 600-1000 ns, Trp-Cage; or 1000-1200 ns, WAAAH-Helix. Across 3 miniproteins, 12 effective temperatures, and 12 solution compositions, we performed ∼ 340 *µ*s of aggregate sampling time. Further details are provided in Tables S1 and S2.

To quantify the solvent effects on protein stability, we define various metrics that capture the protein and solvent as dynamically coupled partners. These metrics are based on quantifying the protein-solvent shape complementarity calculated by representing the protein and solvent networks as three-dimensional shapes. An overview of the approach is shown in Figs. 1 and S6. We considered all protein heavy atoms to build the protein network (PN). All solvent heavy atoms with a minimum distance *r* from the nearest protein heavy atom, such that *r*_*cut*_ ≤ *r* ≤ *r*_*cut*_ + Δ*r*, were used to create the solvent network (SN). Independently, the PN and SN were converted into mesh representations (Fig. S6a) by transforming PN and SN point clouds into convex hulls, with nodes becoming vertices and sets of connected edges forming cycles becoming triangular faces. The protein mesh (PM) and solvent mesh (SM) were “inflated” and “deflated”, respectively, according to the van der Waals radii of the heavy atom corresponding to a given vertex. In this procedure, the position of each vertex was moved radially outwards if the vertex belonged to the PM, or radially inwards for the SM. This process was repeated for configurations extracted from the REST3 trajectories to generate ensembles of PM/SM pairs (Fig. 1a).

**Figure 1.**
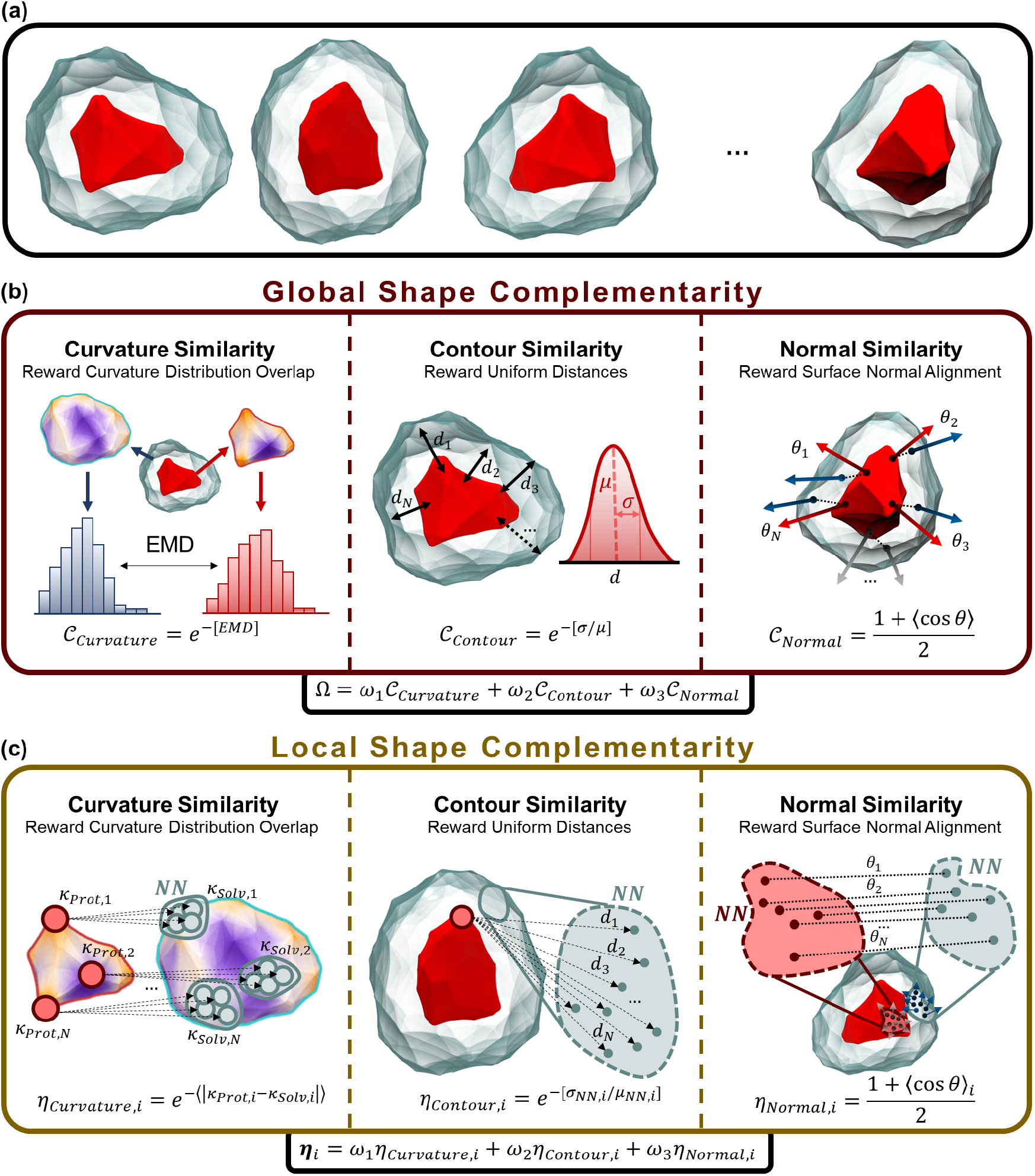
Overview of the protein/solvent shape complementarity scoring functions. (a) Ensemble of protein/solvent shape pairs. (b) Global shape complementarity scores. (c) Local shape complementarity scores. Both global and local scores include measures of curvature similarity, contour similarity, and normal similarity. Red corresponds to protein shapes and points, cyan corresponds to the local solvent network.

Several global (𝒞) and local (*η*) shape similarity metrics were used to assess complementarity between protein and solvent meshes. We derived three similarity metrics (Fig. 1b) – curvature similarity, 𝒞_*Curvature*_; contour similarity, 𝒞_*Contour*_; and normal similarity, 𝒞_*Normal*_. For 𝒞_*Curvature*_, the Cohen-Steiner and Morvan approach^50^ was used to measure the local Gaussian curvature, *κ*, of vertices within the PM and SM. In this method, *κ* is computed for vertex *v* as 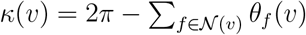, where 𝒩 (*v*) denotes the set of faces incident to vertex *v*, and *θ*_*f*_ (*v*) is the angle at *v* in face *f*. The similarity between probability distributions *P* (*κ*_*PM*_) and *P* (*κ*_*SM*_) was measured by the Earth Mover’s Distance (EMD),^51^ and the resulting score was calculated as 𝒞_*Curvature*_ = ⟨exp[−*EMD*(*P* (*κ*_*PM*_), *P* (*κ*_*SM*_))]⟩, where ⟨·⟩ denotes time-averaged values. The PM/SM pairs with similar curvature distributions (low EMD) have high 𝒞_*Curvature*_ values, indicating complementarity.

Contour similarity, 𝒞_*Contour*_, measures packing inefficiencies between the SM and PM.

_*𝒞Contour*_ was defined as the standard deviation in nearest neighbor protein-solvent distances, *σ*_*NN*_, relative to the average of all nearest neighbor protein-solvent distances, *µ*_*NN*_. Nearest neighbors were identified *via* kd-trees as in Ref. 52. Lower variances indicate an even distribution of protein/solvent vertices and indicate more efficient packing. The time-averaged contour similarity score was calculated as 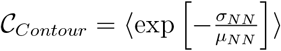. Low variations in protein-solvent distances give higher 𝒞_*Contour*_ values and is associated with high complementarity between PM/SM pairs.

_*𝒞Normal*_ is based on the cos(*θ*) values between normals of the PM and SM triangular faces, measuring the similarity in the orientations of the PM and SM. To calculate 𝒞_*Normal*_, *M* = 1000 points were randomly sampled on the surface of the PM, and for each point, the nearest SM point was identified. The normal of the triangular face containing the PM point was used to calculate a PM vector *u*_*m,P M*_, and similarly, an SM vector *u*_*m,SM*_ was computed. *m* is the index describing the PM or SM point. The average value of cos(*θ*) between normal pairs was computed as 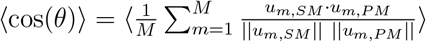. This is then used to compute normal similarity as _*𝒞Normal*_ = (1 + ⟨cos(*θ*) ⟩)*/*2, where values close to 1 represent aligned normals and values near 0 are antiparallel. A composite scoring function was computed as a linear combination of _*𝒞*_ terms, where *ω*_*i*_ represents the weight assigned to similarity measure 𝒞_*i*_: Ω = *ω*_1_ 𝒞 _*Curvature*_ + *ω*_2_ 𝒞 _*Contour*_ + *ω*_3_ 𝒞_*Normal*_.

While _𝒞_ represents global PM/SM shape complementarity measurements, we developed corresponding measurements to assess local, per-residue shape metrics: *η*_*Curvature*_, *η*_*Contour*_, and *η*_*Normal*_ (Fig. 1c). As a local measurement, *η*_*Curvature*_ was based on the absolute difference in local curvatures, *κ*, between PM vertices corresponding to residue *r* and their *k*-nearest SM points: 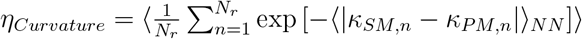, where *N*_*r*_ is the total number of vertices *n* corresponding to residue *r. η*_*Contour*_ was measured as the variation in distances between a PM vertex and the *k*-nearest SM points, 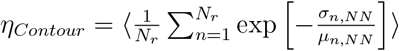, where *NN* denotes an average over *k*-nearest neighbors. *η*_*Normal*_ was taken as angular differences associated with the *k*-nearest PM neighbors of residue vertices *n* and their respective nearest SM point, giving rise to 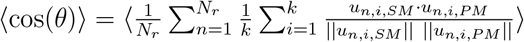 and a score calculation of *η*_*Normal*_ = (1 + ⟨cos(*θ*)⟩)*/*2.

Similar to global similarity metrics, a composite, per-residue scoring function was defined according to a linear combination: ***η*** = *ω*_1_*η*_*Curvature*_ + *ω*_2_*η*_*Contour*_ + *ω*_3_*η*_*Normal*_. To assess whether residue-level shape complementarity is associated with the rank-ordering of excipient effects on temperature stability, the Spearman correlation coefficient, *ρ*_*r*_, between ***η***(*X, r*_*p*_) and Δ*T*_*m*_(*X*) was calculated (*X*: excipient identity; *r*_*p*_ is residue *r* of protein *p*).^53^

The excipient-dependent change in melting temperature (Δ*T*_*m*_) relative to water is shown in Fig. 2, where Δ*T*_*m*_ > 0 indicates greater stability in the excipient solution than in water alone. For Trpzip, the addition of any excipient under study results in improved temperature stability (Δ*T*_*m*_ > 0) (Fig. 2a). Among sugars, the sugar alcohols GOL and SOR are the most effective stabilizers, followed by disaccharides THL and SCRS, while the monosaccharide AGLC is the least effective stabilizer. The most stabilizing amino acid excipients are LEU, ARG, and PHE, all of which feature hydrophobic or partially hydrophobic side chains. In contrast to Trpzip, the majority of excipients destabilize WAAAH-Helix, though ARG, SER, PHE, and GOL have a negligible effect on temperature stability (Fig. 2b). The most destabilizing excipients of WAAAH-Helix include LYS, LEU, and AGLC. Excipients have a wide range of effects on Trp-Cage: ARG and LYS are significantly destabilizing; SER, PHE, LEU, and SCRS are moderately destabilizing; THL and GOL have negligible effects; and GLU, AGLC, and SOR are moderately stabilizing (Fig. 2c). Trp-Cage contains structural elements found in Trpzip and WAAAH-Helix, indicating that the excipient effects may be related to the different structural elements of a given protein.

**Figure 2.**
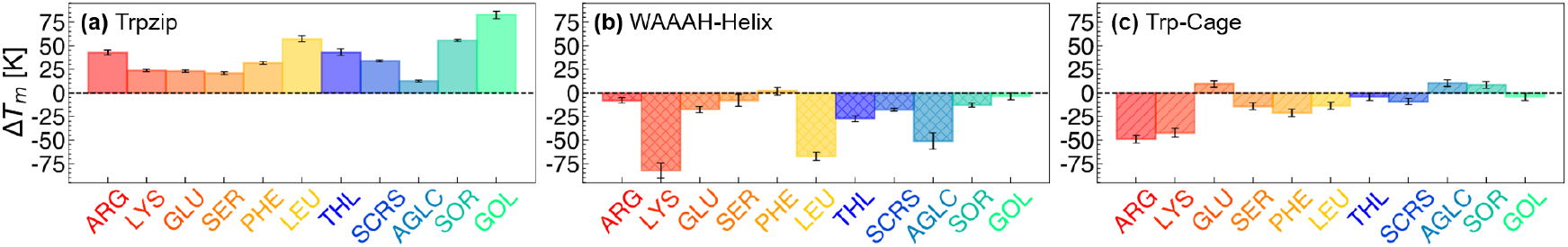
Temperature stability of miniproteins in excipient solutions, as determined from REST3 simulations. (a-c) Change in melting temperature, Δ*T*_*m*_, in excipient solution, relative to water, for (a) Trpzip, (b) WAAAH-Helix, and (c) Trp-Cage.

No single excipient is effective in stabilizing all three miniproteins under study (Fig. 2). This finding alone highlights the significant challenge in formulation design strategies—no universal excipients exist. For certain excipients, such as ARG, we observe a different extent of temperature stabilization for each miniprotein: ARG significantly stabilizes Trpzip, has negligible effect on WAAAH-Helix, and significantly destabilizes Trp-Cage. This finding is supported by previous studies detailing the context-dependent effects of ARG.^31,38,54,55^ We hypothesize that, given ARG stabilizes hydrophobic interactions,^31,38,56^ it is an effective stabilizer of Trpzip, which is primarily stabilized by stacking interactions of dehydrated Trp faces. Because WAAAH-Helix is primarily stabilized by intraprotein hydrogen bonds and not hydrophobic interactions, ARG is ambivalent and does not stabilize or destabilize the helical miniprotein. As for Trp-Cage, we observe competitive ARG interactions with the Arg9-Asp16 salt bridge (Fig. S7), which may significantly disrupt the native state stability of Trp-Cage.

These results demonstrate system-dependent effects of excipients on protein temperature stability. Is there a unified parameter that could explain the broad spectrum of excipient effects? Ideally, such a metric would require no *a priori* knowledge of excipient behavior, instead serving as a predictive tool for determining whether an excipient will stabilize a particular protein. We demonstrate that quantities focused on the relationship between protein and solvent networks can provide such metrics. Specifically, in this paper, we introduce various protein/solvent shape complementarity metrics and demonstrate their correlation with excipient effect on protein stability.

Shape representations (Fig. 1a) of the protein and local solvent environment are built by identifying the smallest convex sets that contain either all protein heavy atoms (protein mesh, PM) or all solvent (water, excipient, and counterions) heavy atoms within a distance from the protein surface (solvent mesh, SM). The similarity of the two shapes are assessed through 𝒞_*Curvature*_, 𝒞_*Normal*_, and 𝒞_*Contour*_. A composite scoring function is obtained as a linear combination of the average values, Ω = Σ_*i*_ *ω*_*i*_C_*i*_, which is used to evaluate the total shape complementarity between the PM and SM. We report results based on weights, *ω*_*i*_, optimized via multivariate regression to predict Δ*T*_*m*_ from Ω (see SI Section 2).

The relationship between Ω and Δ*T*_*m*_ with increasing effective temperatures is shown in Figs. 3a–c and S8. At low temperature replicas, Ω is positively correlated with Δ*T*_*m*_ values of individual proteins (Fig. 3a). All Trpzip/excipient solutions have high Ω values, along with positive Δ*T*_*m*_ values. For WAAAH-Helix, where all excipients either had negligible or destabilizing effects, Ω values are all below 0.5. For Trp-Cage, the stabilizing excipients GLU, AGLC, and SOR have the among highest Ω values. Excipients that significantly destabilize Trp-Cage, such as ARG and LYS, have reduced Ω values. Near the *T*_*m*_ of each protein in water, the relationship between Ω and Δ*T*_*m*_ remains positively correlated (Fig. 3b). Fitting to a sigmoidal function in this effective temperature replica results in *R*^2^ = 0.925, suggesting that shape complementarity captures a substantial fraction of the variance in thermal stabilization near the unfolding transition. Additionally, the dependence of Δ*T*_*m*_ on Ω is nonlinear, with diminishing returns at high shape complementarity values. In other words, modest increases in Ω at low-to-intermediate values correspond to disproportionately large gains in stability, whereas beyond a threshold the stabilizing effect saturates. At the highest temperature replica from REST3 simulations, Ω and Δ*T*_*m*_ relationships become clustered according to protein identity (Fig. 3c). This finding suggests that when proteins are unfolded, the identity of the excipient becomes less important in terms of determining shape complementarity. Alternatively, this may indicate that Ω is dominated by the length of the protein at high effective temperatures (Trpzip > Trp-Cage ≈ WAAAH-Helix).

**Figure 3.**
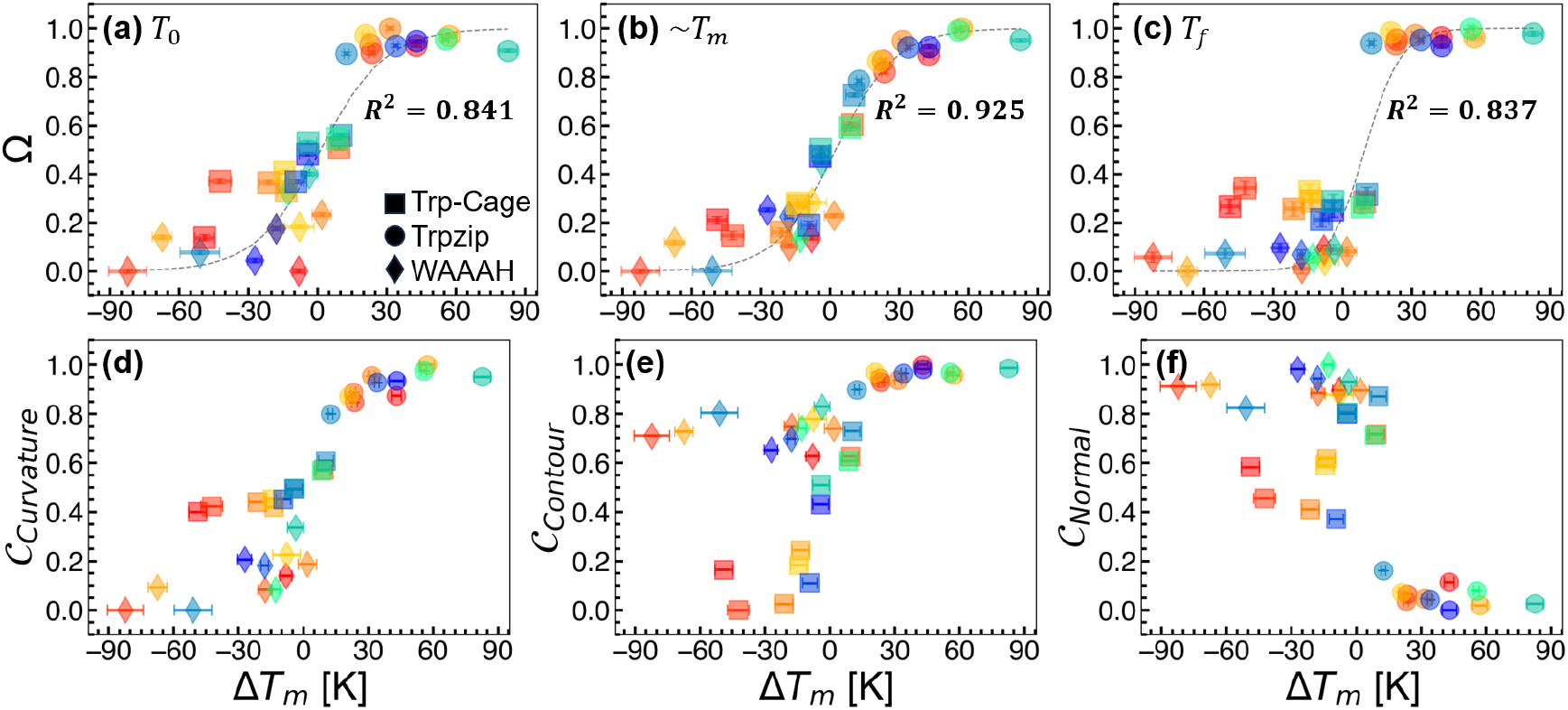
Relationships between protein-solvent shape complementarity and change in the temperature stability of miniproteins. (a-c) Scatter plots showing composite shape complementarity, Ω, against the change in protein melting temperature, Δ*T*_*m*_, following excipient addition. Relationships at different effective temperatures are shown at (a) the lowest temperature replica, *T*_*i*_, (b) the temperature replica closest to the melting temperature in water, *T*_*m*_, and (c) the highest temperature replica, *T*_*f*_. Different proteins are indicated by different shapes (Trp-Cage, square; Trpzip, circle; WAAAH-Helix, diamond), while excipients are indicated by different colors as in Fig. 2. The dashed black line represents a sigmoidal fit in panels (a-c), and corresponding R^2^ values are reported. An *r*_*cut*_ value of 1.0 nm was used to define a Δ*r* = 0.1 nm thick shell of local solvent atoms. (d-f) Scatter plots showing shape complementarity metrics, 𝒞, versus Δ*T*_*m*_. Relationships are shown near *T*_*m*_ for (d) 𝒞_*Curvature*_, (e) 𝒞_*Contour*_, and (f) 𝒞_*Normal*_. Each plot is normalized from 0 to 1.

The relationships between the different C_*i*_ metrics and Δ*T*_*m*_ are shown in Figs. 3d–f and S9–S11. 𝒞_*Curvature*_ and 𝒞_*Contour*_ are positively correlated with Δ*T*_*m*_ (Fig. 3d,e; Tables S3 and S4), while 𝒞_*Normal*_ is negatively correlated with Δ*T*_*m*_ (Fig. 3f; Table S5). High 𝒞_*Curvature*_ (Fig. 3d) values indicate small deviations in the distributions of local Gaussian curvature of PM and SM vertices. The positive correlation between 𝒞_*Curvature*_ and Δ*T*_*m*_ suggests that when the solvent adapts to the curved topology of proteins, solvent-protein interactions are more stabilizing. In general, high values of 𝒞_*Contour*_ (Fig. 3e) correlate with stability. This suggests that uniform solvent packing in the local domain of proteins could contribute as a stabilizing feature. Finally, high 𝒞_*Normal*_ (Fig. 3f) indicates similarity between the orientations of triangular faces of the PM and SM. The negative correlation between 𝒞_*Normal*_ and Δ*T*_*m*_ suggests that, when these normals are too closely aligned, protein-solvent interactions are destabilizing. Normal alignment suggests strongly coupled protein and solvent motions that may restrict orientational degrees of freedom, potentially hindering protein flexibility and thereby, affecting protein stability.

Overall, we observe a positive relationship between protein-solvent shape complementarity and temperature stability. This relationship is consistent across 11 unique excipients and 3 distinct miniproteins, with the strongest relationships observed at the temperature replica closest to the melting temperature of a given miniprotein in water. This indicates that metrics such as shape complementarity can provide a unifying framework to characterize excipient effects on protein stability.

We extend the framework presented to probe how excipients modulate stability at the residue level. To this end, per-residue shape complementarity scores (Figs. S12–S14) are computed, *η*_*Curvature*_, *η*_*Contour*_, and *η*_*Normal*_, and a composite score ***η*** = ∑_*i*_ *ω*_*i*_*η*_*i*_ is calculated. These metrics are closely related to the global shape similarity metrics (Fig. S15). In Fig. 4, per-residue, per-excipient ***η*** values are shown. Data are shown for the temperature replica closest to the miniprotein melting temperature in water, and weights used correspond to the optimized weights identified for global shape complementarity within the same replica (Table S6).

**Figure 4.**
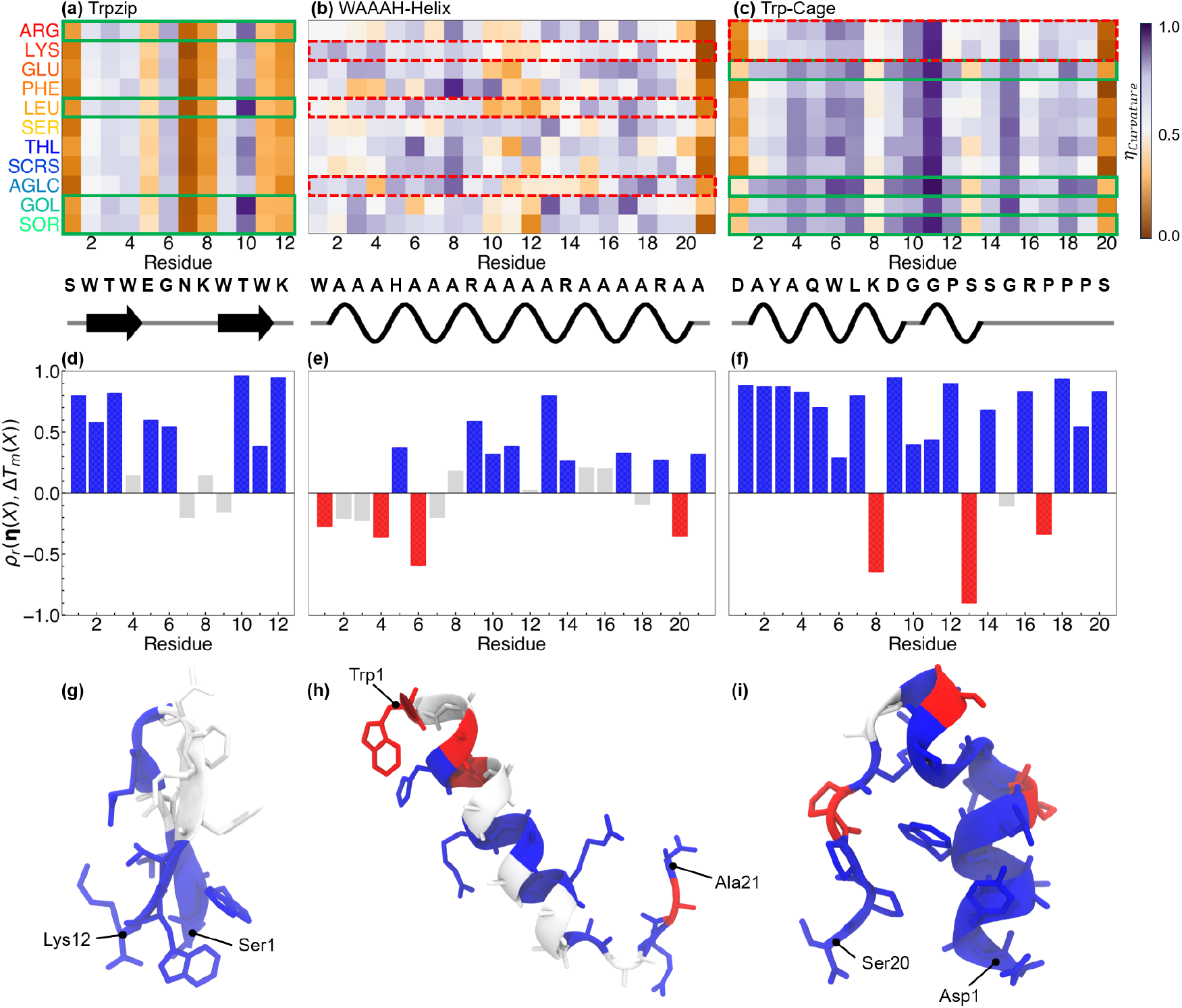
Per-residue shape complementarity scores, *η*. (a-c) Heatmaps are shown for proteins in different excipient solutions, (a) Trpzip, (b) WAAAH-Helix, (c) Trp-Cage. Below each plot, the amino acid sequence is provided according to canonical one-letter codes. Secondary structure assignment was determined by DSSP ^57^ and plotted using Biotite, ^58^ where *β*-sheets are drawn as arrows and *α*-helices are springs. ***η*** is normalized from 0 (orange) to 1 (purple) in (a-c). Green (solid) and red (dashed) boxes are drawn around rows corresponding to stabilizing and destabilizing excipients, respectively. (d-f) Spearman correlation coefficients, *ρ*, of *η* with Δ*T*_*m*_ for (d) Trpzip, (e) WAAAH-Helix, and (f) Trp-Cage. Residues where |*ρ*| < 0.25 are colored gray. (g-i) *ρ* values are projected onto representative configurations of (g) Trpzip, (h) WAAAH-Helix, and (i) Trp-Cage. In (g-i), N- and C-terminal residues are labeled. In (d-i), Colors represent negative correlation (red) and positive correlation (blue).

***η*** represents the local protein-solvent shape complementarity in the vicinity of individual residues. In Trpzip, local shape complementarity is moderate near residues Trp2, Thr3, Trp4, Gly6, and Trp9, and most prominent near Thr10 (Fig. 4a). Conversely, low shape complementarity is observed in the vicinity of Trp11, terminal residues Ser1 and Lys12, as well as loop-forming residues Glu5, Asn7, and Lys8. These observations are broadly true for all excipients, though some variation across excipients exists. In WAAAH-Helix, high complementarity residues (regardless of excipient identity) include His5, Ala8, Arg9, Ala16, and Ala17 (Fig. 4b). Less complementary networks are observed for residues Ala10, Ala11, and Ala12, as well as for the terminal Ala21 residue. Interestingly, when compared to Trp-Cage and Trpzip, the variability across different excipient solutions is markedly more prominent in WAAH-Helix. For Trp-Cage (Fig. 4c), complementary solvent networks are observed around residues Ala4, Gln5, Trp6, Leu7, Asp9, Gly10, Pro12, Gly15, Pro17, Pro18, and Pro19, and most significantly for Gly11, for all excipients under study. Less complementary residues include Asp1, Lys8, Ser13, and Ser20. Across all three miniproteins and 11 excipients, the following trends emerge: (i) less complementary networks tend to form around lysine, serine, and terminal residues, and (ii) more complementary networks tend to form around glycine, proline, and tryptophan residues. Given the high propensity for glycine and proline to appear in loops and turns,^59,60^ these results suggest that secondary structure context may contribute to protein-solvent shape complementarity. This is further supported by the poor complementarity observed for terminal residues, which are frequently more flexible and located outside of regular secondary structural elements.

In Fig. 4d-f, we plot *ρ*_*r*_(*η*(*X*), Δ*T*_*m*_(*X*)), where positive values indicate the surrounding solvent environment around a given residue is more complementary in stabilizers, while negative values indicate the local shape complementarity is poor in stabilizers. In other words, this analysis enables the identification of residues where mismatches in protein-solvent shape complementarity may be *beneficial* to stability. In Trpzip, Trp-Trp stacking interactions and Thr-Thr hydrogen-bonding are among the most crucial interactions in the folding process.^39,61,62^ Potentially, complementary protein-solvent networks near these residues promote intraprotein interactions among tryptophan residues, improving stability. We observed a strong positive correlation between shape complementarity and change in melting temperature for two of the core Trp residues, Trp2 and Trp11, the hydrogen-bond “zipper” forming residues Thr3 and Thr10, and terminal residues Ser1 and Lys12 (Fig. 4d). Indeed, the most effective Trpzip stabilizers, ARG, LEU, GOL, and SOR, are also the excipients that promote the highest protein-solvent shape complementarity around these residues.

WAAAH-Helix is primarily stabilized *via* intrahelical hydrogen bonds, ^48,63^ though it has been shown that increased Trp1-His5 distance is also associated with improved native fold stability.^64^ Interestingly, high shape complementarity around the key His5 residue is correlated with stability, while high complementarity around Trp1 is associated with destabilization (Fig. 4e). We hypothesize that mismatched Trp1–His5 complementarity prevents these residues from forming contacts that promote off-pathway, misfolded states. Dissimilar solvent networks around Ala20, which is situated near the C-terminus, are also associated with increased temperature stability. The most prominent WAAAH-Helix destabilizers, LYS, LEU, and AGLC, notably have complementarity profiles that vary from one another.

Key Trp-Cage residues involved in stabilization of the folded state include Pro17, Pro18, and Pro19, which form a hydrophobic core alongside Tyr3, Trp6, and Leu7.^41,43,65^ Residues 1–9 form an N-terminal helix, and Gly10, Gly11, and Pro12 form the middle 3_10_-helix. Stabilizing excipients for Trp-Cage, including GLU, AGLC, and GOL, form more complementary solvent networks around Leu7, Asp9, and Pro18 relative to other excipients (Fig. 4c). In Trp-Cage, most residues have a positive correlation between ***η*** and Δ*T*_*m*_ (Fig. 4f). Low complementarity for residues Lys8, Ser13, and Pro17 is also correlated with increased temperature stability. These findings highlight the observed diversity in excipient effects on the stability of the miniproteins. While there are some general trends, the subtle interplay of various interactions modulates the effects observed. This further emphasizes the need for the global and local analysis presented here.

Given the correlation between protein-solvent shape complementarity in the miniproteins across 33 different solutions, we evaluated if such a framework also extends to larger globular proteins. We studied hen egg white lysozyme (HEWL) and thaumatin (TMT) in excipient solutions (Fig. 5c). The melting temperatures were estimated from differential scanning fluorimetry (DSF) experiments (Table S8).^66,67^ Protein-solvent shape complementarity was calculated from a collective 1.2 *µ*s NPT simulations per protein/excipient pair in 0.1 M excipient solutions (ARG, LYS, GLU, THL, SCRS, and SOR). These solutions were simulated near the experimental melting temperatures of each protein in water (337 K for TMT; 343 K for HEWL) for 12 independent 100 ns simulations (Fig. S16– S19). We observed, however, that our results are consistent even if only sampling near the native state is considered (Fig. S20).

**Figure 5.**
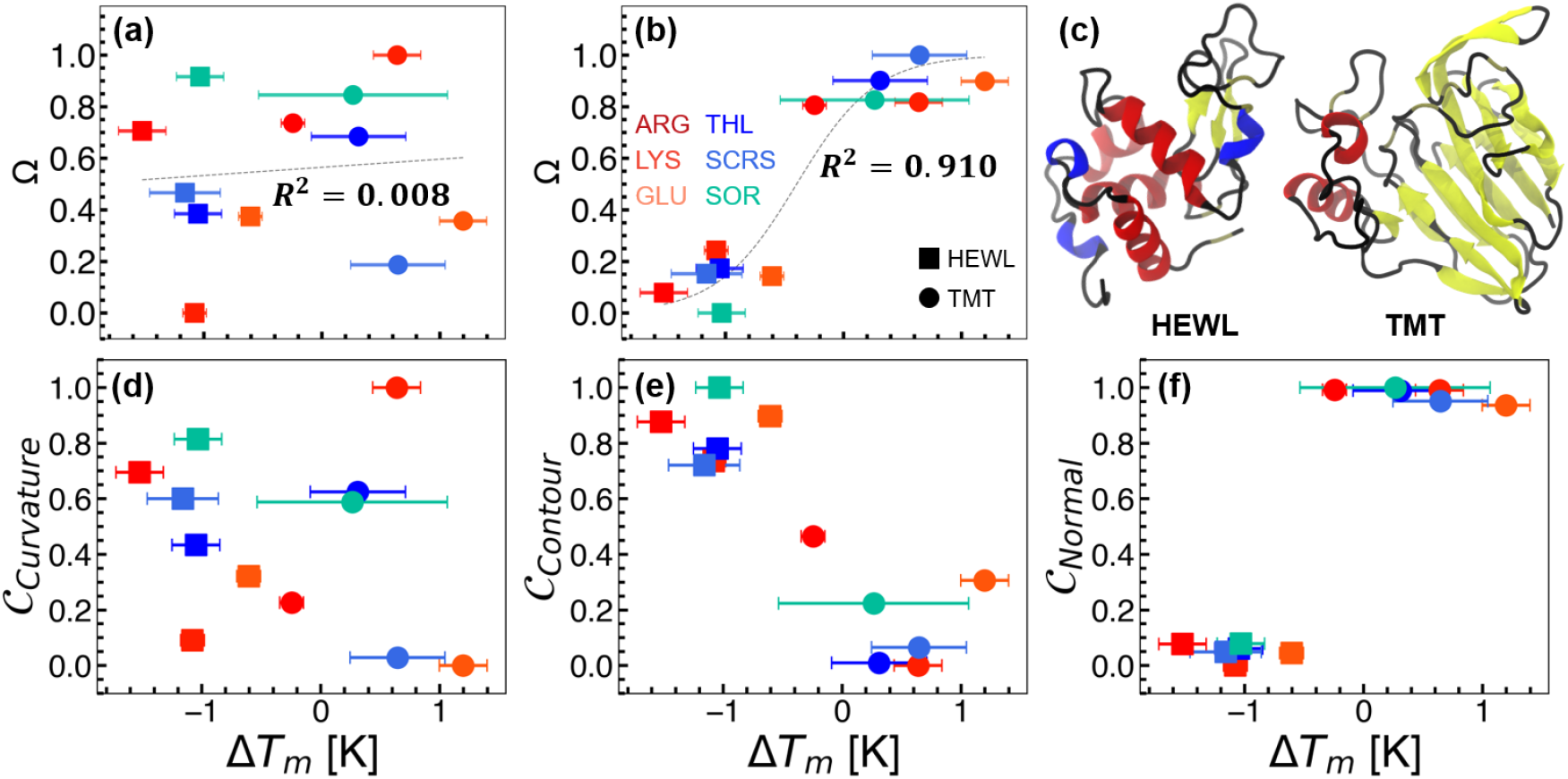
Relationships between protein-solvent shape complementarity and change in the temperature stability of proteins. (a,b) Scatter plots showing composite shape complementarity, Ω, against the change in protein melting temperature, Δ*T*_*m*_, as determined experimentally. Ω is computed using (a) the same weights as determined for miniproteins and (b) weights obtained *via* linear regression on all excipients. Different proteins are indicated by different shapes (HEWL, square; TMT, circle), while excipients are indicated by different colors as in Fig. 2. The dashed black line represents a sigmoidal fit and the corresponding R^2^ value is shown. Error bars represent the standard deviation from three replicate samples. (c) Representative structures of globular proteins under study. (d-f) Scatter plots of shape complementarity values versus Δ*T*_*m*_, (d) 𝒞_*Curvature*_, (e) 𝒞_*Contour*_, and (f) 𝒞_*Normal*_.

We first evaluated whether the weights obtained for the various contributions to Ω obtained from miniproteins were transferable to the globular protein systems. Fig. 5a shows that the correlation between the Ω and Δ*T*_*m*_ is poor. This result indicates that, relative to miniproteins, the same weights from 𝒞_*Curvature*_, 𝒞_*Contour*_, and 𝒞_*Normal*_ are not sufficient to predict excipient-mediated stability. To examine this further, multiple linear regression was carried out to optimize *ω*_*i*_ for protein/excipient combinations.

Optimal *ω*_*i*_ values were obtained through multiple linear regression using data from all excipients (Fig. 5b), only amino acids (Fig. S21a) and only sugars (Fig. S21e). In all cases, we observe a strong correlation between Ω and Δ*T*_*m*_. This indicates that even for larger, globular proteins, a relationship exists between the local solvent environment and protein stability. Interestingly, the weight of each contributing term to the shape complementarity varies between the proteins and miniproteins (Table S7).

We find that curvature similarity and contour similarity are negatively correlated with protein stability, whereas normal similarity is positively correlated (Fig. 5d-f). This is different from the observations in miniproteins, where curvature and contour similarity were positively correlated with stability, while a negative correlation between normal similarity with stability was observed. Several factors could contribute to this difference – in the case of miniproteins, we sample the folded-to-unfolded configurational space, while for the globular proteins, our conformational sampling is limited. Additionally, the structure, shape, and flexibility of globular proteins differ from those of miniproteins. Thus, while the signs and magnitudes of individual correlations may depend on the sampled conformational ensemble and protein class, the observation that protein–solvent shape complementarity correlates with excipient-dependent stability in both miniproteins and globular proteins supports the transferability of our framework.

In the design of biological formulations, the mechanisms of the stabilizing or destabilizing effects of excipients on biomolecular stability are often unknown. Recent work has focused on shifting trial-and-error excipient selection to a more information-driven approach, ^10–13,31,32,68^ which depends on understanding how excipients interact with proteins at the molecular level. Excipient effects are highly system-dependent, however, due to the chemical complexity and diversity of proteins. Adding to this challenge is the fact that no single physical property is predictive of excipient effectiveness.

In this paper, we demonstrate that a single physical property, protein-solvent shape complementarity, has the potential to predict excipient-mediated changes in miniprotein melting temperatures. Specifically, stabilizing excipients form complementary solvent networks in the local domain of three miniproteins with diverse structural features and folding mechanisms. Destabilizing excipients, meanwhile, form less complementary networks around these proteins. Furthermore, we demonstrate that the framework is transferable to globular proteins. Our findings are well aligned with the perspective that shifts mechanistic explanations for cosolvent-mediated stability away from traditional theories centered on preferential interactions or exclusion,^69,70^ and toward an emphasis on the hydration shell as the focal point. Recent work suggests that similarity between water structure in osmolyte and protein solutions provides a metric for classifying osmolytes: stabilizing osmolytes have hydration spheres that are structurally compatible with those of the protein, whereas denaturing osmolytes have structurally distinct hydration spheres.^19–21,23^ Another body of work focuses on osmolyte-mediated modulation of protein-associated water dynamics.^18,24–27,33,34^ Stabilizing osmolytes reduce the translational and rotational dynamics of water in the protein hydration shell, while denaturants increase hydration shell dynamics. The shape complementarity framework presented here unifies these perspectives by quantifying the ensemble-averaged structural compatibility between proteins and their surrounding solvent, thereby implicitly incorporating the underlying solution dynamics.

Our framework extends well beyond excipient and co-solute screening. While the importance of solvent effects on protein stability and function is well established, most analyses treat protein structure and hydration water in isolation. In contrast, our framework explicitly couples these contributions, accounting for both protein and solvent structure and dynamics in a unified description. This positions it as a general tool for interrogating how the solvent environment shapes protein behavior, applicable wherever the interplay between a protein and its surroundings is of mechanistic interest.

Overall, our findings highlight the power of molecular simulations to transform formulation science from empirical guesswork into a predictive discipline. By revealing that shape complementarity between proteins and the local solvent environment can be manipulated by excipient selection, this work lays the foundation for rational excipient design. As the chemical space of biological therapeutics continues to expand, such mechanistic insights are indispensable for designing formulations that reliably preserve protein integrity across diverse therapeutic and industrial applications. More broadly, this framework connects microscopic solvent organization to macroscopic thermodynamic behavior and applies to a wide range of problems involving biomolecular assembly, conformational transitions, and molecular recognition, where the solvent plays a critical role in governing dynamics and function.

## Author Contributions

J.W.P.Z.: Investigation, Conceptualization, Validation, Formal Analysis, Visualization, Writing - Original Draft, Writing - Review and Editing; P.M.: Validation, Writing - Review and Editing; X.Z.: Investigation - Experimental; C.L.H.: Funding Acquisition, Writing - Review and Editing; S.L.P.: Funding Acquisition, Writing - Review and Editing; S.S.: Conceptualization, Methodology, Supervision, Project Administration, Funding Acquisition, Resources, Writing - Review and Editing

## Data availability

Data for this article, including simulation parameter files and analysis scripts, are available at https://github.com/SAMPEL-Group/Miniproteins-Excipients.

## Acknowledgments

This work is supported by U.S. National Science Foundation DMREF grants #2118788, #2118693, and #2325392. Computational resources were provided by the Minnesota Super-computing Institute at the University of Minnesota - Twin Cities. J. W. P. Z. acknowledges funding support from the University of Minnesota Graduate School Doctoral Dissertation Fellowship. S. L. P. and X. Z. thank Prof. Michelle Farkas in the Department of Chemistry at UMass Amherst for the use of the PCR instrument for differential scanning fluorimetry experiments.

## Supporting information

Supporting information: details on convergence checks, error estimations, additional computational details, and additional experimental details.

