## Supporting Information for "Protein-Solvent Shape Complementarity as a Unifying Principle in Excipient-Mediated Protein Thermal Stability"

### Contents


**1 Simulation Protocols** **S-3**

1.1 System Set-up and Simulation Parameters . . . . . . . . . . . . . . . . . . . . S-3

1.2 Replica Exchange with Solute Tempering 3 (REST3) Simulations . . . . . . S-4

**2 Mesh Construction** **S-12**

2.1 Overview . . . . . . . . . . . . . . . . . . . . . . . . . . . . . . . . . . . . . . S-12

2.2 Global Shape Complementarity . . . . . . . . . . . . . . . . . . . . . . . . . S-15

2.3 Local Shape Complementarity . . . . . . . . . . . . . . . . . . . . . . . . . . S-22

2.4 Normalization and Error Estimates . . . . . . . . . . . . . . . . . . . . . . . S-27

2.5 Linear Regression and Sigmoidal Fine-Tuning . . . . . . . . . . . . . . . . . S-28

2.6 Extension to Globular Proteins . . . . . . . . . . . . . . . . . . . . . . . . . S-29

2.7 Pseudocodes . . . . . . . . . . . . . . . . . . . . . . . . . . . . . . . . . . . . S-35

**3 Experimental Temperature Stability Assays** **S-37**

3.1 Materials . . . . . . . . . . . . . . . . . . . . . . . . . . . . . . . . . . . . . S-37

3.2 Stock Solution Preparation . . . . . . . . . . . . . . . . . . . . . . . . . . . . S-37

3.3 Sample Preparation . . . . . . . . . . . . . . . . . . . . . . . . . . . . . . . . S-37

3.4 Thermal Shift Characterization . . . . . . . . . . . . . . . . . . . . . . . . . S-38

**References** **S-40**

### 1 Simulation Protocols

#### 1.1 System Set-up and Simulation Parameters

All simulations were performed using GROMACS 2021.4[S1,S2] with the PLUMED 2.8.0[S3,S4] patch applied. Systems were initially subject to energy minimization using the steepest descent algorithm.[S5] An NVT equilibration was carried out for 1 ns at 300 K using the V-rescale thermostat[S6] followed by a 1 ns NPT equilibration using the stochastic cell rescaling barostat.[S7] During equilibration, we used coupling constants of $\tau_T = 0.1$ ps$^{-1}$ and $\tau_P = 0.5$ ps$^{-1}$ for the thermostat and barostat, respectively. Following equilibration, NPT production runs were completed using the Nosé-Hoover thermostat[S8] and Parrinello-Rahman barostat,[S9] with coupling constants of $\tau_T = 5$ ps$^{-1}$ for the protein, $\tau_T = 1$ ps$^{-1}$ for the solvent, and $\tau_P =$ 10 ps$^{-1}$. The Particle Mesh Ewald (PME) algorithm[S10] was used for electrostatic interactions with a cut-off of 1.2 nm. A reciprocal grid of 42 x 42 x 42 cells was used with 4$^{\text{th}}$ order B-spline interpolation. A force-switching scheme from 1–1.2 nm was used to describe long-range van der Waals interactions. The neighbor search was performed every 10 steps. Lorentz-Berthelot mixing rules[S11,S12] were used to calculate non-bonded interactions between different atom types. The CHARMM36 force field[S13,S14] was used to parameterize protein and amino acid excipient molecules. For sugar excipients, we used a modified CHARMM parameter set optimized to reproduce sugar Kirkwood-Buff integrals.[S15,S16] All systems were solvated with TIP3P water.[S17] Experimentally-resolved structures of Trpzip (Trpzip1 variant; PDB: 1LE0),[S18] Trp-Cage (tc10b variant; PDB: 2JOF),[S19] Thaumatin (from *Thaumatococcus daniellii*; PDB: 4BAL)[S20] and Lysozyme (from hen egg white; PDB: 2YVB) were used to model proteins. WAAAH-Helix was initially modeled using the Avogadro software[S21] by threading the protein sequence onto an $\alpha$-helical backbone. The protonation states of titratable protein residues at pH 7 were determined by the PROPKA3 method.[S22] For each system, excipient molecules were added to reach an initial concentration of 0.1 M. $Na^+$ and $Cl^-$ atoms were added to neutralize each system, as necessary.

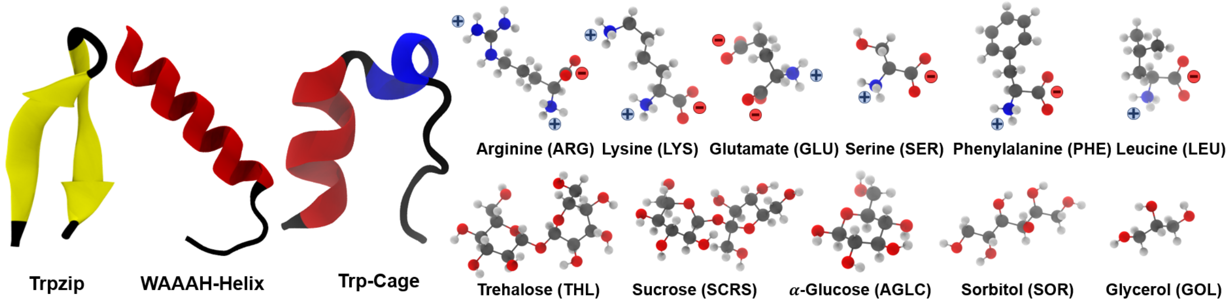


Figure S1: Selected miniprotein and excipient molecules used in the present study. Miniproteins are shown in cartoon representation and are colored according to secondary structure ($\beta$-strand – yellow; $\alpha$-helix – red; $3_{10}$-helix – blue; coils/turns - black). Excipients are shown in ball-and-stick representation and are colored according to atom type (C – gray; O – red; N – blue; H – white). Positive and negative charge groups are denoted by '+' and '-' symbols, respectively.

#### 1.2 Replica Exchange with Solute Tempering 3 (REST3) Simulations

To efficiently explore the conformational landscapes of miniprotein folding/unfolding, we utilized Replica Exchange with Solute Tempering 3 (REST3) protocol.[S23] REST3 builds on the original REST and REST2 protocols,[S24,S25] which use Hamiltonian rescaling to simulate different regions of the system under different effective temperatures. The scaled Hamiltonian at condition $m$ is given as:

$$E_m^{REST} = \lambda_m^{PP} E_{PP}(X) + \lambda_m^{PW} E_{PW}(X) + \lambda_m^{WW} E_{WW}(X) \tag{S1}$$

where $X$ denotes the system coordinates, $\lambda$ is a scaling factor, and the energy of the system is decomposed into solute-solute $E_{PP}$, solute-solvent $E_{PW}$, and solvent-solvent $E_{WW}$ energies. The manner in which the scaling factors are determined is the difference between REST, REST2, and REST3. In the original REST protocol:

$$\lambda_m^{PP} = \beta_m/\beta_0,\ \lambda_m^{pw} = (\beta_0 + \beta_m)/2\beta_0,\ \lambda_m^{ww} = 1 \tag{S2}$$

where $\beta_m = 1/k_B T_m$, $\beta_0 = 1/k_B T_0$, and $k_B$ is the Boltzmann constant. In REST2:

$$\lambda_m^{PP} = \beta_m/\beta_0,\ \lambda_m^{pw} = \sqrt{\beta_m/\beta_0},\ \lambda_m^{ww} = 1 \tag{S3}$$

which weakened solute-solvent interactions compared to the original REST protocol, alleviating certain bottlenecks between low- and high-temperature conditions. This, however, leads to over-compaction of proteins at high temperatures.[S23] REST3 recovers proper chain expansion by treating solute-solvent van der Waals (vdW) interactions as an independent, tunable parameter:

$$E_m^{REST3} = \lambda_m^{PP} E_{PP}(X) + \lambda_m^{PW} E_{PW}^{elec} + \kappa_m \lambda_m^{PW} E_{PW}^{vdW}(X) + \lambda_m^{WW} E_{WW}(X) \tag{S4}$$

where $\kappa_m$ is an additional scaling factor for solute-solvent vdW interactions at condition $m$. For Trp-Cage and Trpzip, we applied a linear relationship to set $\kappa_m$ at different effective temperatures:

$$\kappa_m = \begin{cases} 1, & m < 4 \\ 1 + 0.005 * (m - 3), & m \geq 4 \end{cases} \tag{S5}$$

To recover the experimental $T_m$ of WAAAH-Helix, we found it was necessary to weaken protein-solvent interactions at lower temperatures (Fig. S2). We were able to achieve this by setting $\kappa_m$ according to:

$$\kappa_m = 1 + 0.025 * (m - 12) \tag{S6}$$

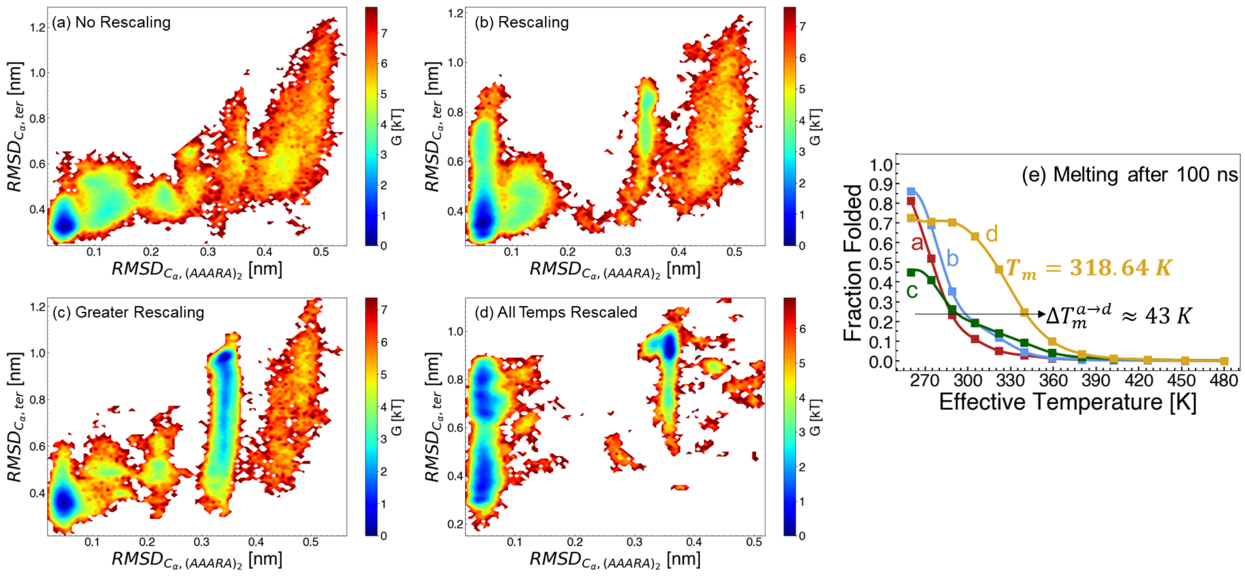


Figure S2: Change in WAAAH-Helix free energy landscapes with different protein-water interaction scaling strengths. (a) No rescaling, $\kappa = 1.0, 1.0, 1.0, 1.0, 1.0, 1.005..., 1.035$, (b) Rescaling of all replicas with effective temperature less than 300 K, $\kappa = 0.98, 0.985, ..., 1.035$ (c) Stronger rescaling of replicas with effective temperature less than 300 K, $\kappa = 0.94, 0.945, ..., 1.025$, and (d) Rescaling of all temperature replicas, $\kappa = 0.7, 0.725, ..., 0.975$

Table S1: Miniprotein/excipient systems under study. Numbers shown include the total number of atoms ($N$), the number of water molecules ($N_{WAT}$), excipient molecules ($N_{Exc}$), sodium ions ($N_{Na^+}$), and chloride ions ($N_{Cl^-}$).

| **System** | $N$ | $N_{WAT}$ | $N_{Exc}$ | $N_{Na^+}$ | $N_{Cl^-}$ |
|---|---|---|---|---|---|
| Trp-Cage / WAT | 13565 | 4427 | 0 | 0 | 0 |
| Trp-Cage / ARG | 13526 | 4330 | 9 | 0 | 9 |
| Trp-Cage / LYS | 13529 | 4337 | 9 | 0 | 9 |
| Trp-Cage / GLU | 13517 | 4354 | 9 | 9 | 0 |
| Trp-Cage / SER | 13553 | 4381 | 9 | 0 | 0 |
| Trp-Cage / PHE | 13508 | 4339 | 9 | 0 | 0 |
| Trp-Cage / LEU | 13532 | 4350 | 9 | 0 | 0 |
| Trp-Cage / THL | 13514 | 4275 | 9 | 0 | 0 |
| Trp-Cage / SCRS | 13556 | 4289 | 9 | 0 | 0 |
| Trp-Cage / AGLC | 13544 | 4348 | 9 | 0 | 0 |
| Trp-Cage / SOR | 13532 | 4338 | 9 | 0 | 0 |
| Trp-Cage / GOL | 13550 | 4380 | 9 | 0 | 0 |
| Trpzip / WAT | 14541 | 4774 | 0 | 0 | 1 |
| Trpzip / ARG | 14505 | 4678 | 9 | 0 | 10 |
| Trpzip / LYS | 14547 | 4698 | 9 | 0 | 10 |
| Trpzip / GLU | 14494 | 4702 | 9 | 8 | 0 |
| Trpzip / SER | 14529 | 4728 | 9 | 0 | 1 |
| Trpzip / PHE | 14526 | 4700 | 9 | 0 | 1 |
| Trpzip / LEU | 14520 | 4701 | 9 | 0 | 1 |
| Trpzip / THL | 14499 | 4636 | 9 | 0 | 1 |
| Trpzip / SCRS | 14529 | 4635 | 9 | 0 | 1 |
| Trpzip / AGLC | 14511 | 4692 | 9 | 0 | 1 |
| Trpzip / SOR | 14529 | 4692 | 9 | 0 | 1 |
| Trpzip / GOL | 14550 | 4735 | 9 | 0 | 1 |
| WAAAH-Helix / WAT | 26778 | 8833 | 0 | 0 | 3 |
| WAAAH-Helix / ARG | 26733 | 8734 | 9 | 0 | 12 |
| WAAAH-Helix / LYS | 26733 | 8740 | 9 | 0 | 12 |
| WAAAH-Helix / GLU | 26736 | 8762 | 9 | 6 | 0 |
| WAAAH-Helix / SER | 26742 | 8779 | 9 | 0 | 3 |
| WAAAH-Helix / PHE | 26748 | 8754 | 9 | 0 | 3 |
| WAAAH-Helix / LEU | 26766 | 8763 | 9 | 0 | 3 |
| WAAAH-Helix / THL | 26775 | 8697 | 9 | 0 | 3 |
| WAAAH-Helix / SCRS | 26730 | 8682 | 9 | 0 | 3 |
| WAAAH-Helix / AGLC | 26760 | 8755 | 9 | 0 | 3 |
| WAAAH-Helix / SOR | 26751 | 8746 | 9 | 0 | 3 |
| WAAAH-Helix / GOL | 26733 | 8776 | 9 | 0 | 3 |

Table S2: REST3 simulation lengths, per replica. Simulation times depended on both the identity of the protein and the absence/presence of excipient molecules.

| **Miniprotein** | **Excipients?** | **Simulation Time per Replica [ns]** | **Total Time Over 12 Replicas [$\mu$s]** | **Total Time Over All Systems [$\mu$s]** |
|---|---|---|---|---|
| Trpzip | No | 200 | 2.4 | 2.4 |
| WAAAH-Helix | No | 1200 | 14.4 | 14.4 |
| Trp-Cage | No | 600 | 7.2 | 7.2 |
| Trpzip | Yes | 400 | 4.8 | 52.8 |
| WAAAH-Helix | Yes | 1000 | 12.0 | 132.0 |
| Trp-Cage | Yes | 1000 | 12.0 | 132.0 |

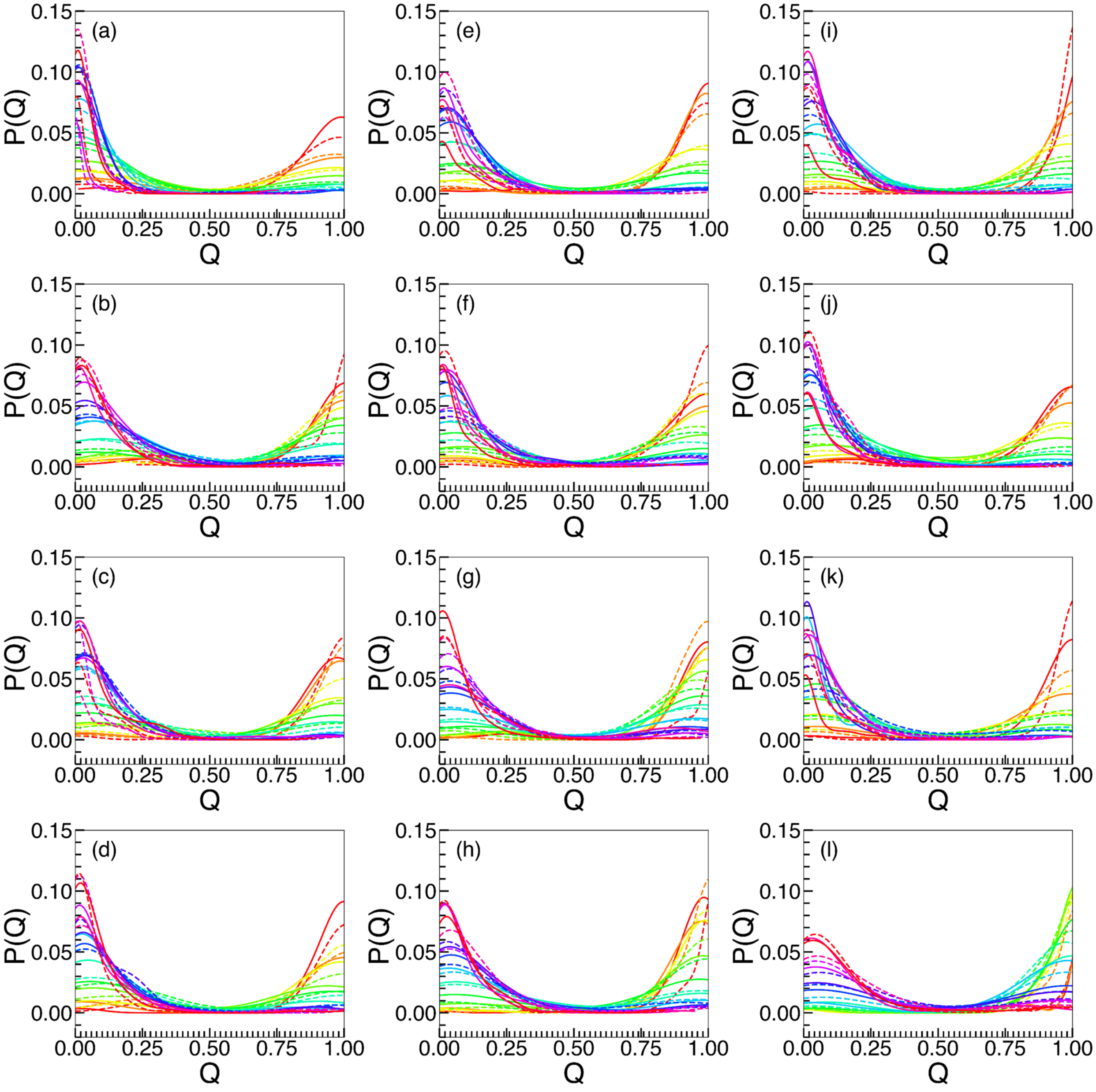


Figure S3: Probability distributions of the fraction of native contacts for Trpzip. Data are shown for two equal blocks of time (dashed and solid lines). Colors indicate different replicas. (a) Water, (b) ARG, (c) LYS, (d) GLU, (e) SER, (f) PHE, (g) LEU, (h) THL, (i) SCRS, (j) AGLC, (k) SOR, (l) GOL.

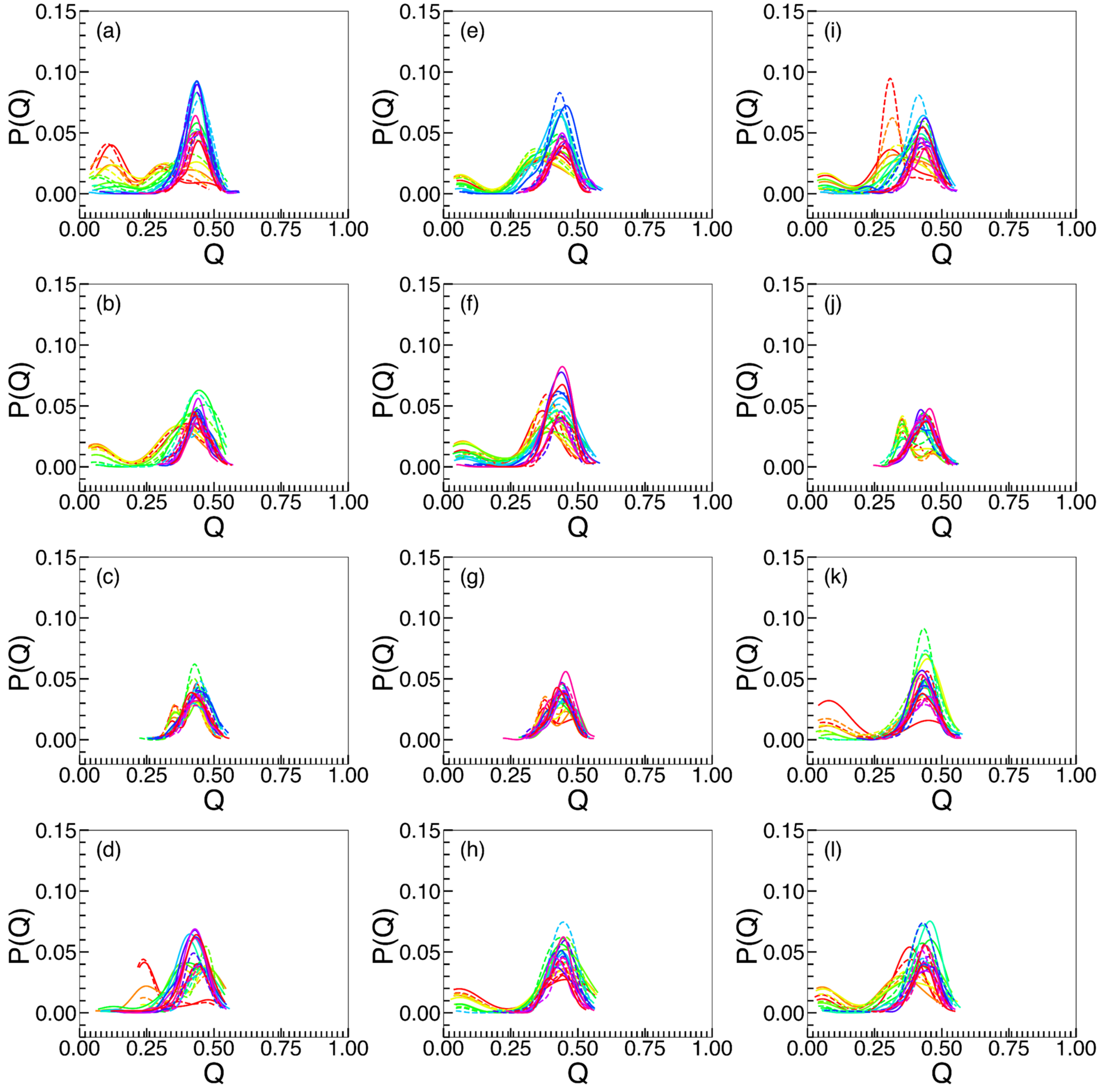


Figure S4: Probability distributions of the fraction of native contacts for WAAAH-Helix. Data are shown for two equal blocks of time (dashed and solid lines). Colors indicate different replicas. (a) Water, (b) ARG, (c) LYS, (d) GLU, (e) SER, (f) PHE, (g) LEU, (h) THL, (i) SCRS, (j) AGLC, (k) SOR, (l) GOL.

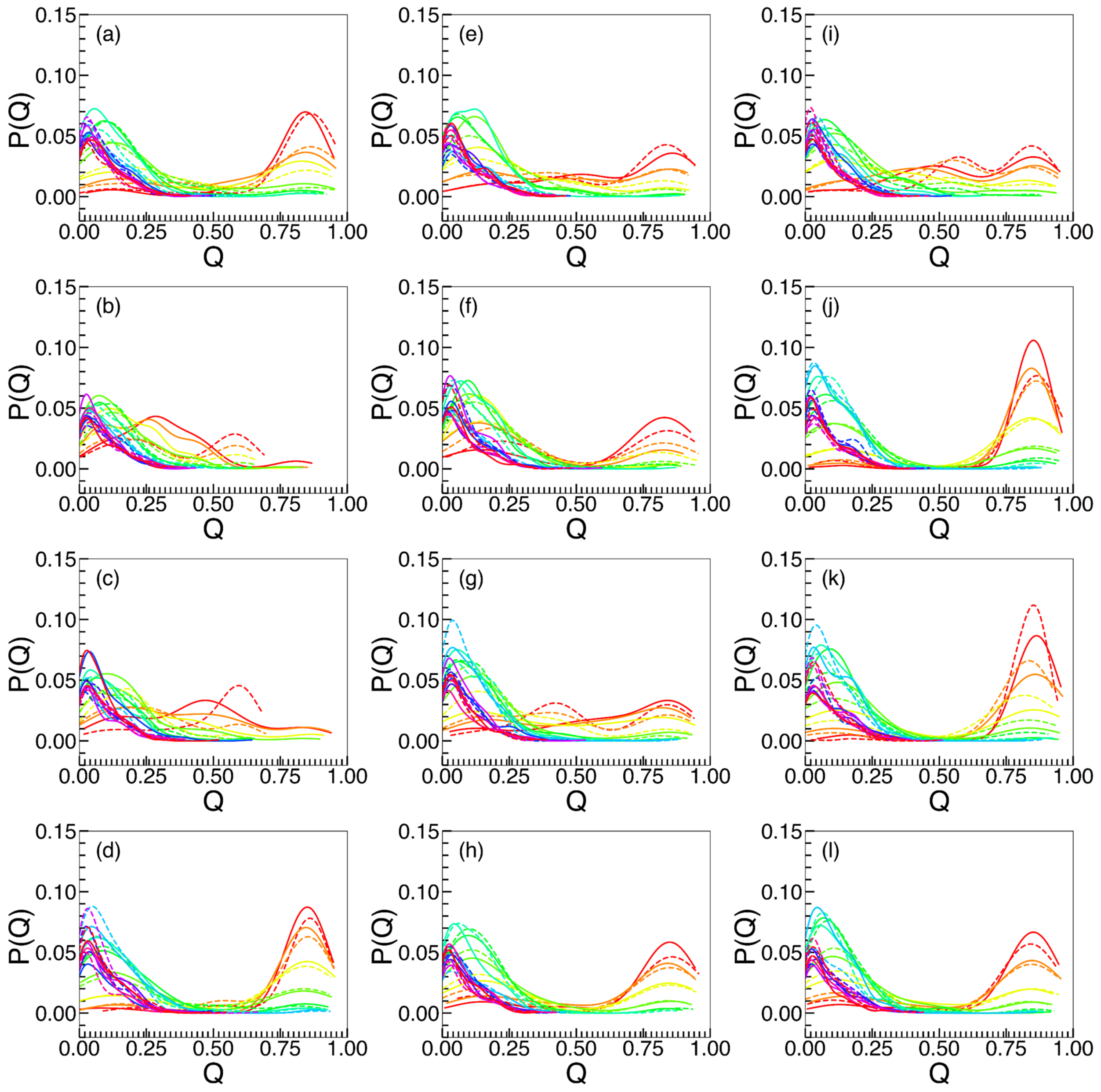

Figure S5: Probability distributions of the fraction of native contacts for Trp-Cage. Data are shown for two equal blocks of time (dashed and solid lines). Colors indicate different replicas. (a) Water, (b) ARG, (c) LYS, (d) GLU, (e) SER, (f) PHE, (g) LEU, (h) THL, (i) SCRS, (j) AGLC, (k) SOR, (l) GOL.

### 2 Mesh Construction

#### 2.1 Overview

All protein heavy atoms were assigned to the protein network (PN). Solvent heavy atoms whose minimum heavy-atom distance to any protein heavy atom was less than the cutoff distance $r_{cut}$ were assigned to the solvent network (SN). Distances were computed using the minimum-image convention under periodic boundary conditions. PN and SN were treated independently in all subsequent geometric procedures. An overview of mesh construction and scoring is shown in Fig. S6.

Convex hulls of PN and SN were constructed using the Quickhull algorithm.[S26] This procedure yielded a watertight, globally convex triangular mesh used for quantities such as enclosed volume and surface normals. The convex hull was computed using the standard Quickhull implementation in SciPy. Interior facets generated by coplanar point sets were removed using the library's default facet-merging routines. The output consisted of a triangular mesh whose vertices corresponded to a subset of the original atom positions.

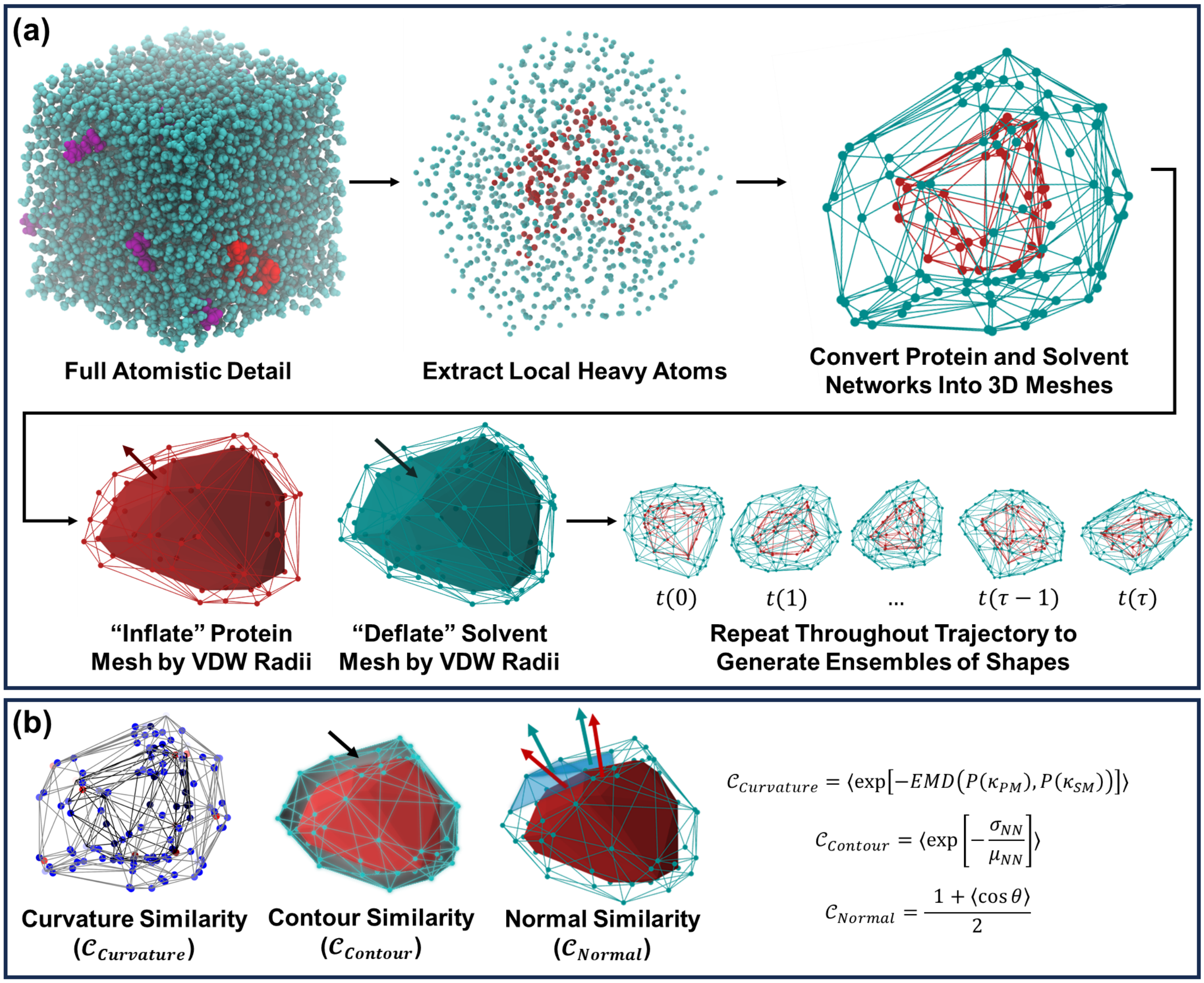


Figure S6: Overview of the protein/solvent shape complementarity analysis workflow. (a) Shape construction workflow. Protein atoms and meshes are red, while solvent atoms and meshes are cyan. (b) Shape similarity metrics, $\mathcal{C}_{Curvature}$, $\mathcal{C}_{Normal}$, and $\mathcal{C}_{Contour}$.

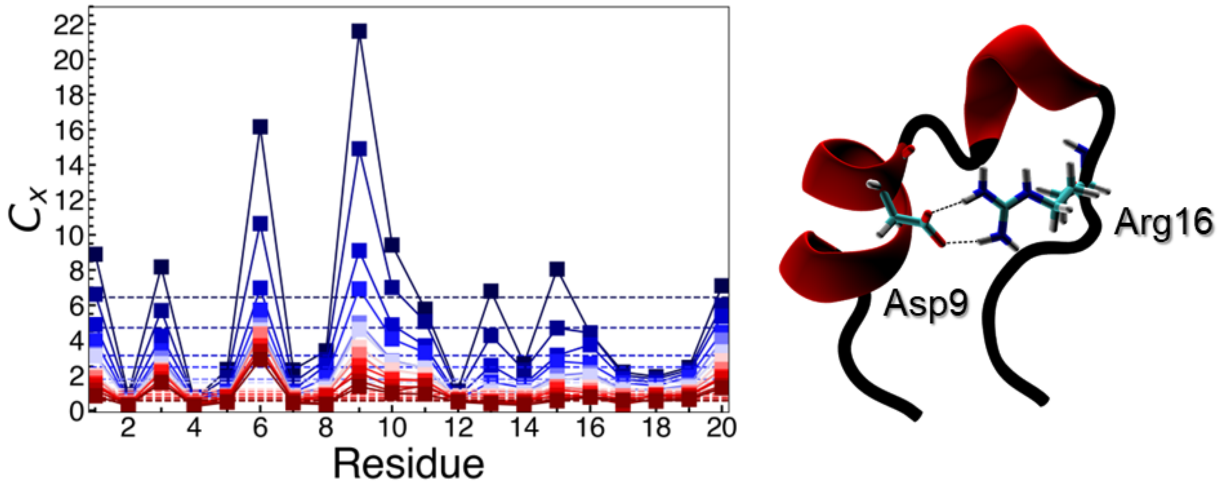


Figure S7: Contact coefficient ($C_x$) associated with ARG and Trp-Cage. $C_x = \left(\frac{N_{Exc}}{N_{Wat}}\right)\left(\frac{M_{Wat}}{M_{Exc}}\right)$, where $N$ represents the local number of excipient (Exc) or water (Wat) atoms, and $M$ represents the total number of Exc or Wat atoms in the system.

#### 2.2 Global Shape Complementarity

Composite shape scores at each effective temperature are provided in Fig. S8. The weights obtained following multivariate regression in each subplot in Fig. S8 are provided in Table S6. Similarly, relationships between $\mathcal{C}_i$ and $\Delta T_m$ are provided in Figs. S9– S11. Spearman rank correlation coefficients corresponding to these figures are provided in Tables S3–S5.

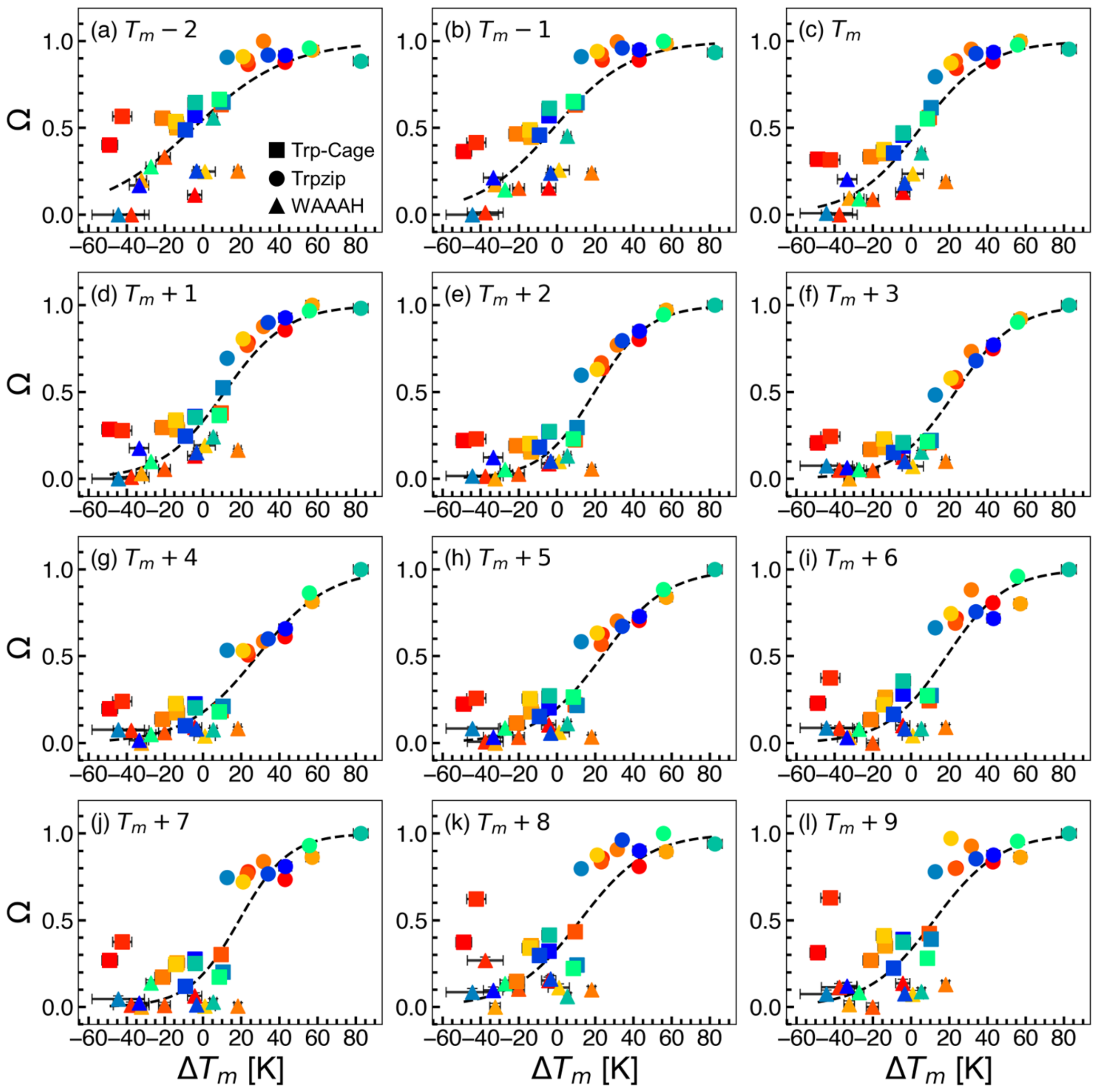


Figure S8: Scatter plots showing composite shape complementarity ($\Omega$) against the change in protein melting temperature ($\Delta T_m$), as determined by REST3 simulations, following excipient addition. Relationships at different effective temperatures are shown. Plot labels indicate the number of replicas away the effective temperature is from the wild-type melting temperature. Different proteins are indicated by different shapes (Trp-Cage, square; Trpzip, circle; WAAAH, triangle), while excipients are indicated by the coloring scheme provided in Fig. 2.

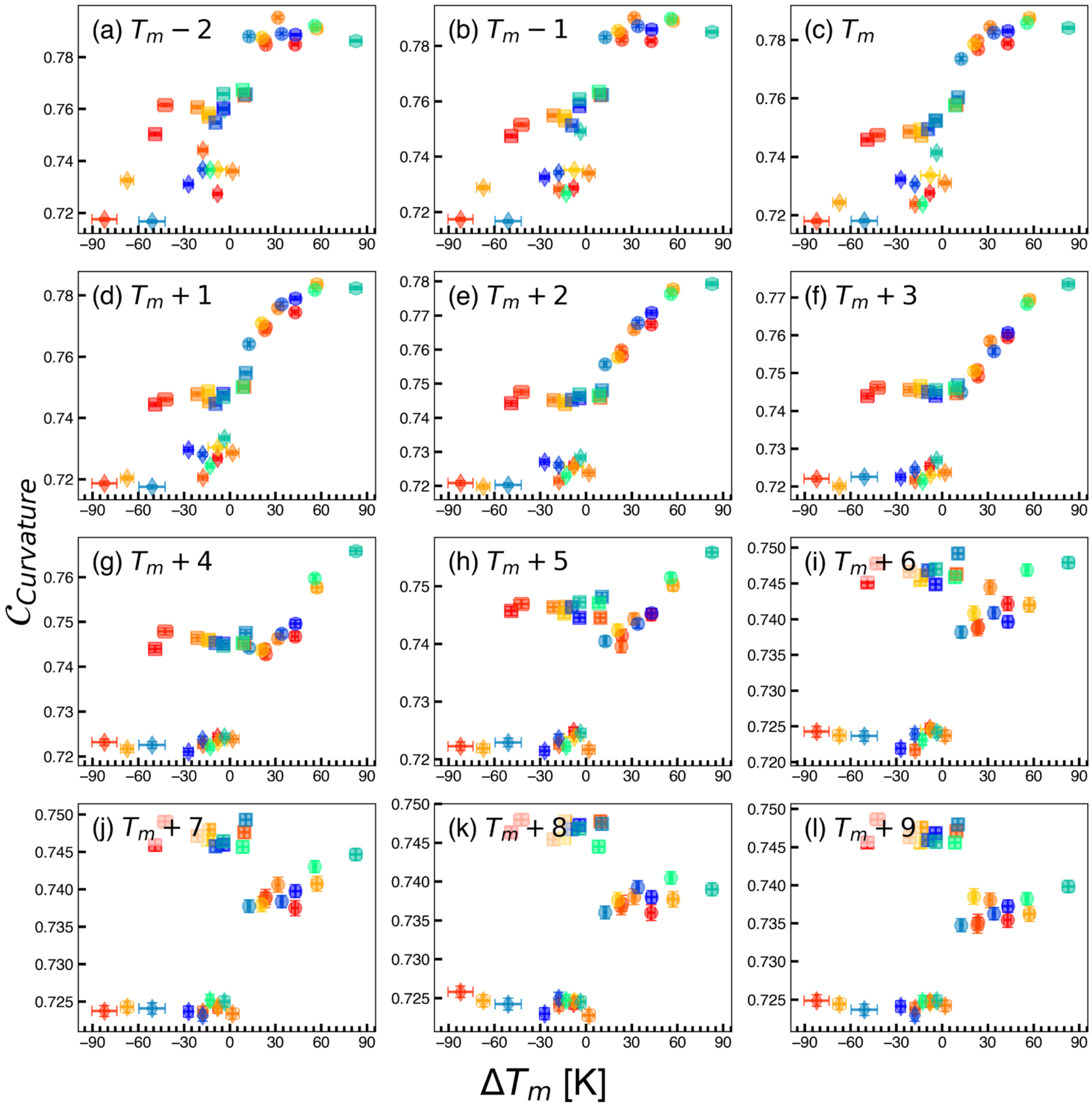


Figure S9: Change in $\mathcal{C}_{Curvature}$ shape similarity metrics in different REST3 replicas. Plot labels indicate the number of replicas away the effective temperature is from the wild-type melting temperature. Different proteins are indicated by different shapes (Trp-Cage, square; Trpzip, circle; WAAAH, triangle), while excipients are indicated by the coloring scheme provided in Fig. 2.

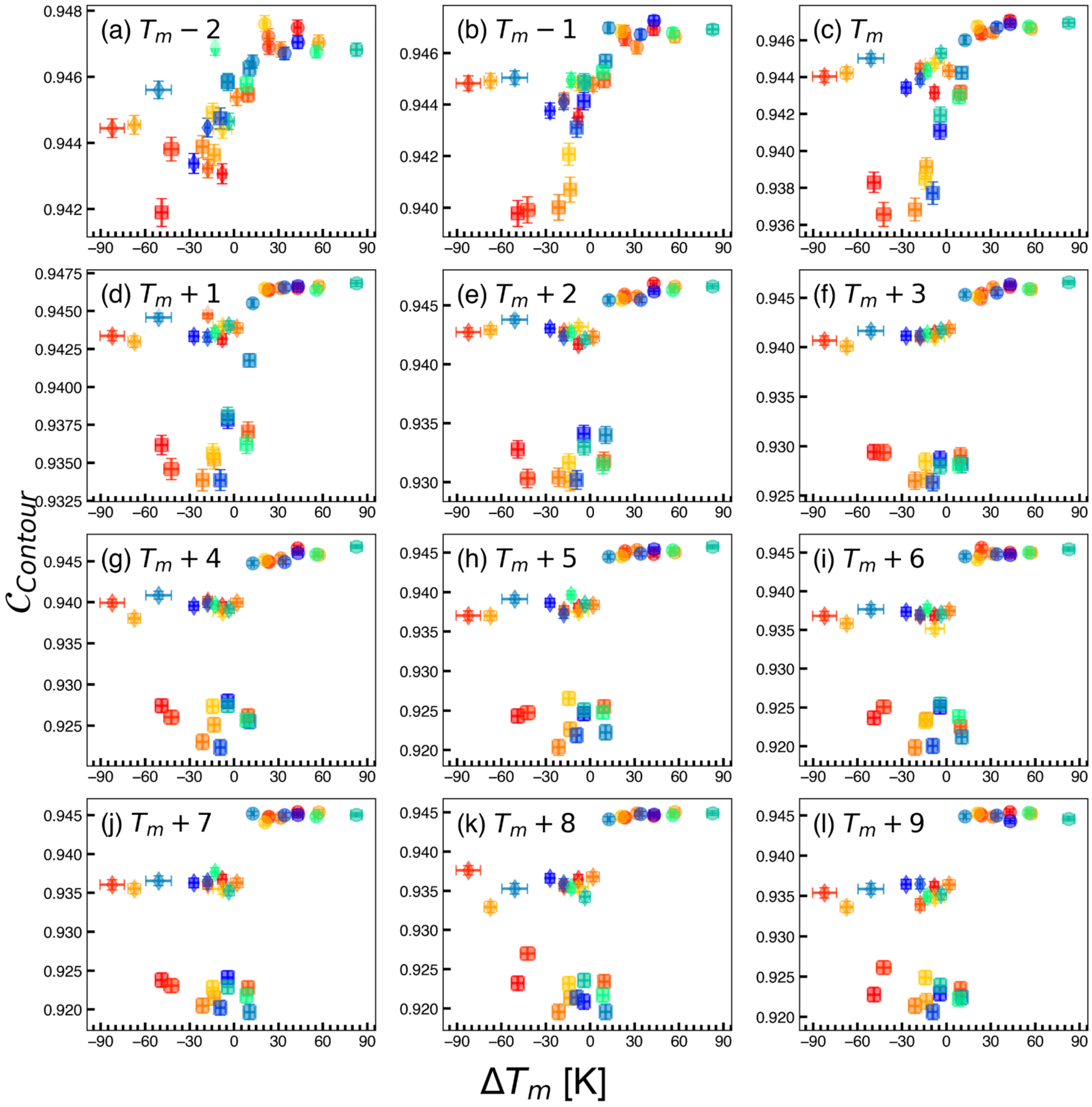


Figure S10: Change in $\mathcal{C}_{Contour}$ shape similarity metrics in different REST3 replicas. Plot labels indicate the number of replicas away the effective temperature is from the wild-type melting temperature. Different proteins are indicated by different shapes (Trp-Cage, square; Trpzip, circle; WAAAH, triangle), while excipients are indicated by the coloring scheme provided in Fig. 2.

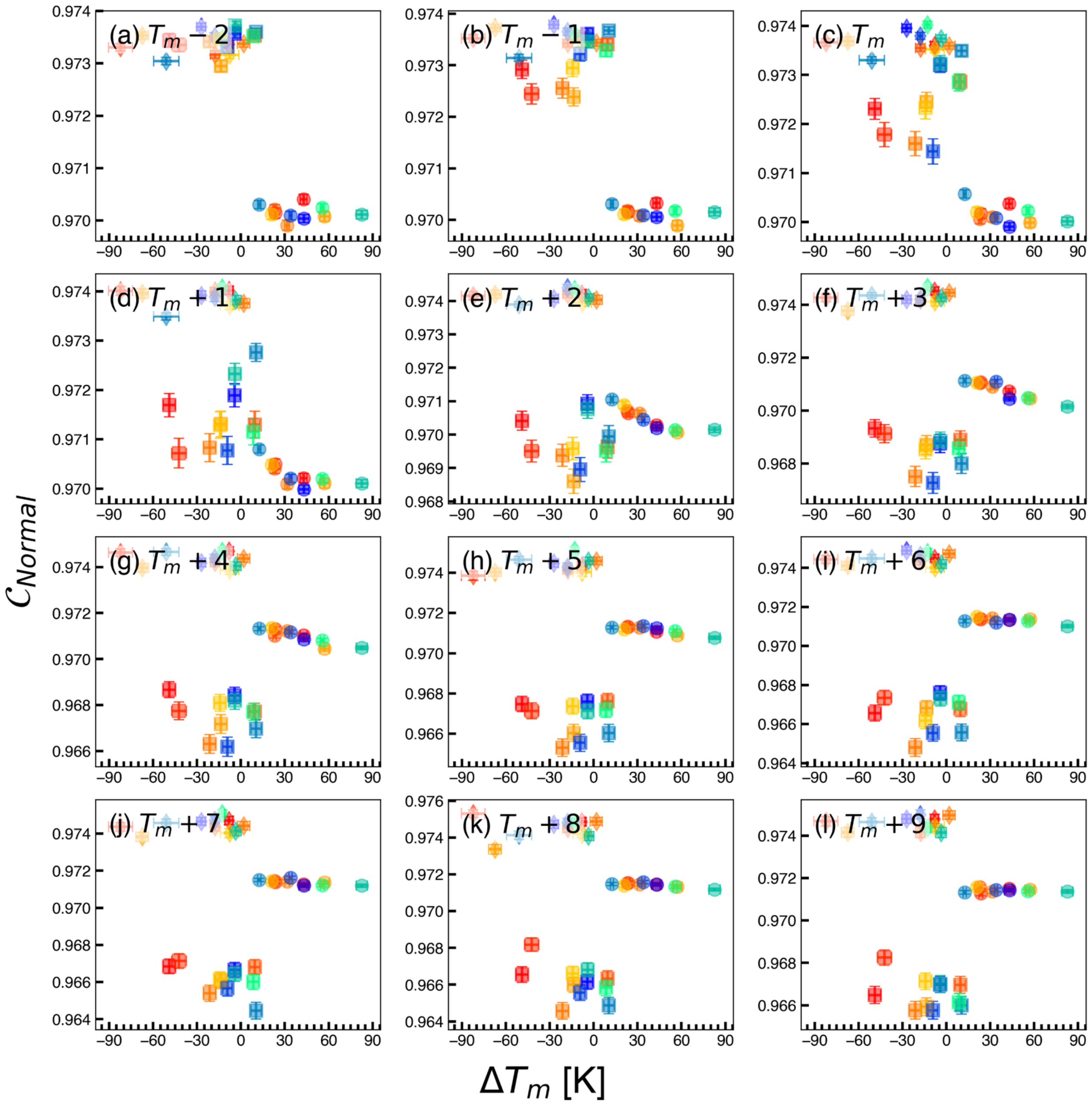


Figure S11: Change in $\mathcal{C}_{Normal}$ shape similarity metrics in different REST3 replicas. Plot labels indicate the number of replicas away the effective temperature is from the wild-type melting temperature. Different proteins are indicated by different shapes (Trp-Cage, square; Trpzip, circle; WAAAH, triangle), while excipients are indicated by the coloring scheme provided in Fig. 2.

Table S3: Spearman correlation coefficients ($\rho$) from different REST3 indices (i) in the comparison of $\mathcal{C}_{Curvature}$ and $\Delta T_m$ following excipient addition. $\rho$ values are provided for comparisons of all proteins (Total), Trpzip (TZ), WAAAH-Helix (WH), and Trp-Cage (TC).

| i | $\rho_{\text{Total}}$ | $\rho_{\text{TZ}}$ | $\rho_{\text{WH}}$ | $\rho_{\text{TC}}$ |
|---|---|---|---|---|
| 0 | 0.8440 | 0.2000 | 0.5550 | 0.6640 |
| 1 | 0.8620 | 0.3640 | 0.6090 | 0.8450 |
| 2 | 0.8830 | 0.7910 | 0.6730 | 0.9450 |
| 3 | 0.8790 | 0.9360 | 0.7360 | 0.7360 |
| 4 | 0.8550 | 0.9820 | 0.5640 | 0.5640 |
| 5 | 0.8010 | 0.9640 | 0.6000 | 0.1640 |
| 6 | 0.6240 | 0.9000 | 0.6640 | -0.0730 |
| 7 | 0.4420 | 0.9360 | 0.3180 | 0.2270 |
| 8 | 0.3170 | 0.7550 | 0.1820 | 0.2550 |
| 9 | 0.1820 | 0.7180 | 0.1640 | 0.0640 |
| 10 | 0.1440 | 0.5640 | -0.4000 | 0.1090 |
| 11 | 0.1800 | 0.4820 | 0.1550 | 0.1180 |

Table S4: Spearman correlation coefficients ($\rho$) from different REST3 indices (i) in the comparison of $\mathcal{C}_{Contour}$ and $\Delta T_m$ following excipient addition. $\rho$ values are provided for comparisons of all proteins (Total), Trpzip (TZ), WAAAH-Helix (WH), and Trp-Cage (TC).

| i | $\rho_{\text{Total}}$ | $\rho_{\text{TZ}}$ | $\rho_{\text{WH}}$ | $\rho_{\text{TC}}$ |
|---|---|---|---|---|
| 0 | 0.7920 | -0.0910 | 0.0640 | 0.8450 |
| 1 | 0.7890 | -0.0270 | -0.2180 | 0.9820 |
| 2 | 0.7290 | 0.6450 | 0.3000 | 0.9000 |
| 3 | 0.6920 | 0.7450 | 0.3180 | 0.6820 |
| 4 | 0.6150 | 0.7910 | -0.5180 | 0.4090 |
| 5 | 0.6570 | 0.7910 | 0.6360 | -0.1910 |
| 6 | 0.6150 | 0.7820 | -0.1550 | 0.0000 |
| 7 | 0.6630 | 0.7450 | 0.3360 | 0.1910 |
| 8 | 0.6270 | 0.5910 | 0.1820 | -0.0730 |
| 9 | 0.5830 | 0.4730 | -0.0910 | -0.3820 |
| 10 | 0.6110 | 0.8000 | 0.0000 | -0.2360 |
| 11 | 0.5960 | -0.0450 | 0.0730 | -0.0820 |

Table S5: Spearman correlation coefficients ($\rho$) from different REST3 indices (i) in the comparison of $\mathcal{C}_{Normal}$ and $\Delta T_m$ following excipient addition. $\rho$ values are provided for comparisons of all proteins (Total), Trpzip (TZ), WAAAH-Helix (WH), and Trp-Cage (TC).

| i | $\rho_{\text{Total}}$ | $\rho_{\text{TZ}}$ | $\rho_{\text{WH}}$ | $\rho_{\text{TC}}$ |
|---|---|---|---|---|
| 0 | -0.5690 | -0.3090 | 0.0360 | 0.6550 |
| 1 | -0.6340 | -0.2550 | -0.2180 | 0.8000 |
| 2 | -0.7000 | -0.5180 | -0.1000 | 0.7550 |
| 3 | -0.7370 | -0.7450 | -0.2270 | 0.4550 |
| 4 | -0.2860 | -0.9730 | -0.1360 | 0.2820 |
| 5 | -0.1630 | -0.8550 | 0.3550 | -0.2360 |
| 6 | -0.1970 | -0.8910 | -0.1180 | -0.2450 |
| 7 | -0.1330 | -0.6450 | 0.3550 | 0.0730 |
| 8 | -0.1520 | -0.5360 | 0.0270 | 0.0820 |
| 9 | -0.1930 | -0.7820 | 0.0550 | -0.3820 |
| 10 | -0.1790 | -0.6450 | 0.0820 | -0.3090 |
| 11 | -0.1550 | -0.0550 | -0.0450 | -0.1270 |

#### 2.3 Local Shape Complementarity

Local shape complementarity scores at the REST3 replica closest to $T_m$ are provided in Figs. S12– S14. The correlation between averaged local and global shape complementarity scores are provided in Fig. S15. A strong, positive correlation is observed for each comparison, indicating that local and global measurements are capturing related shape features.

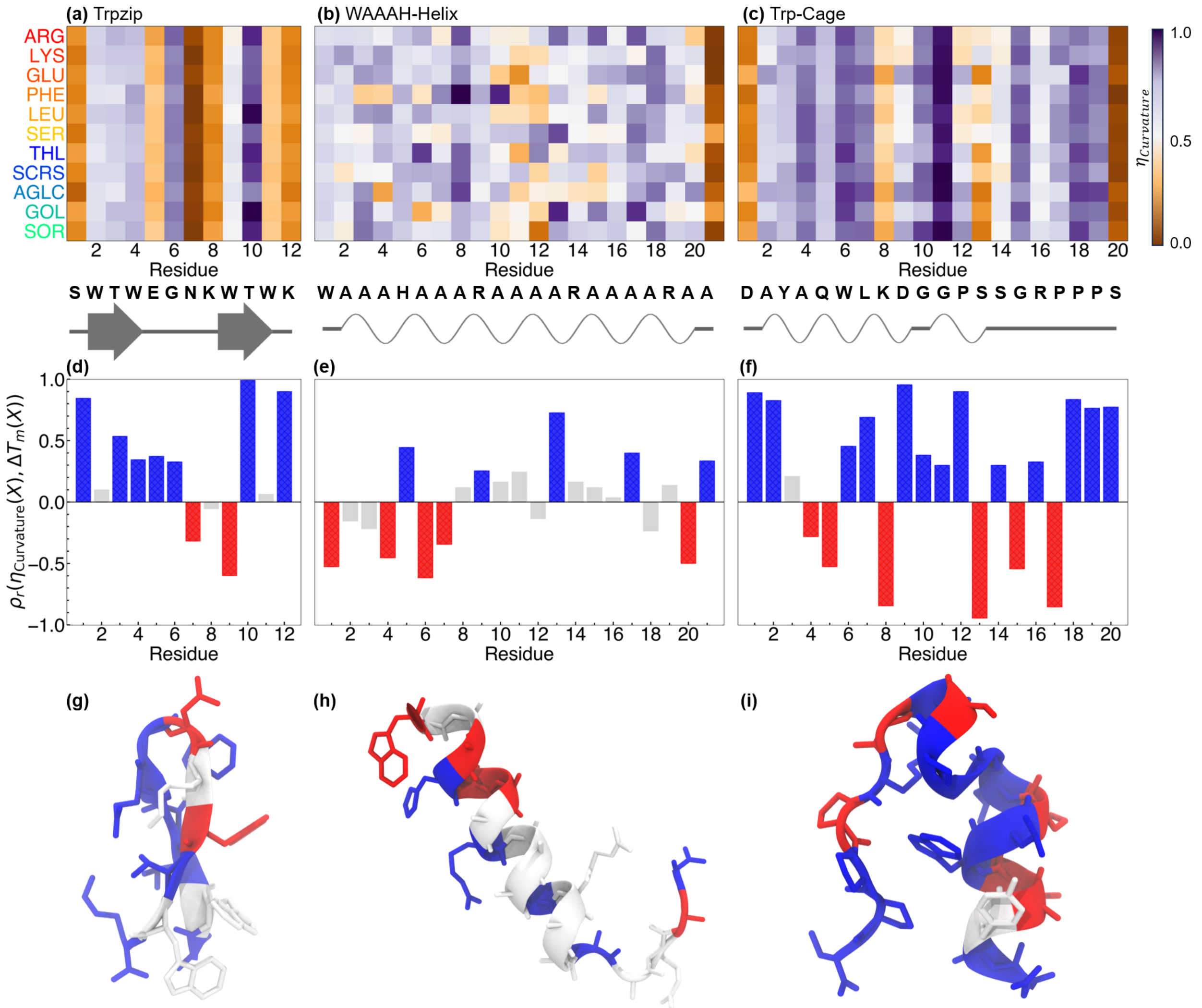


Figure S12: Per-residue curvature similarity scores, $\eta_{Curvature}$. (a-c) Heatmaps are shown for proteins in different excipient solutions, (a) Trpzip, (b) WAAAH-Helix, (c) Trp-Cage. Below each plot, the amino acid sequence is provided according to canonical one-letter codes. Secondary structure assignment was determined by DSSP[S27] and plotted using Biotite,[S28] where $\beta$-sheets are drawn as arrows and $\alpha$-helices are springs. $\boldsymbol{\eta}$ is normalized from 0 (orange) to 1 (purple) in (a-c). (d-f) Spearman correlation coefficients, $\rho$, of $\eta$ with $\Delta T_m$ for (d) Trpzip, (e) WAAAH-Helix, and (f) Trp-Cage. Colors represent negative correlation (red) and positive correlation (blue). Residues where $|\rho| < 0.25$ are colored gray. (g-i) $\rho$ values are projected onto representative configurations of (g) Trpzip, (h) WAAAH-Helix, and (i) Trp-Cage.

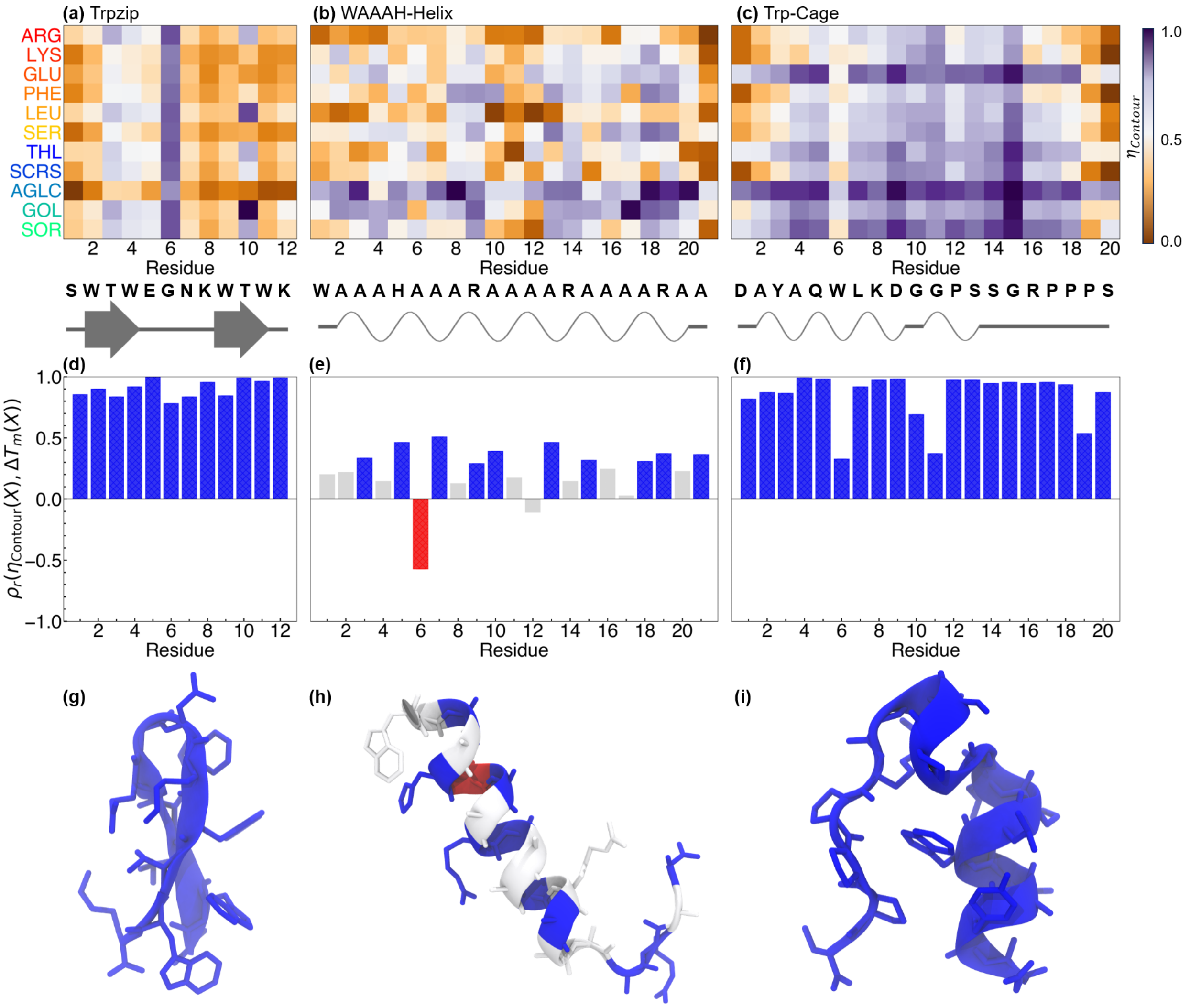


Figure S13: Per-residue contour similarity scores, $\eta_{Contour}$. (a-c) Heatmaps are shown for proteins in different excipient solutions, (a) Trpzip, (b) WAAAH-Helix, (c) Trp-Cage. Below each plot, the amino acid sequence is provided according to canonical one-letter codes. Secondary structure assignment was determined by DSSP[S27] and plotted using Biotite,[S28] where $\beta$-sheets are drawn as arrows and $\alpha$-helices are springs. $\boldsymbol{\eta}$ is normalized from 0 (orange) to 1 (purple) in (a-c). (d-f) Spearman correlation coefficients, $\rho$, of $\eta$ with $\Delta T_m$ for (d) Trpzip, (e) WAAAH-Helix, and (f) Trp-Cage. Colors represent negative correlation (red) and positive correlation (blue). Residues where $|\rho| < 0.25$ are colored gray. (g-i) $\rho$ values are projected onto representative configurations of (g) Trpzip, (h) WAAAH-Helix, and (i) Trp-Cage.

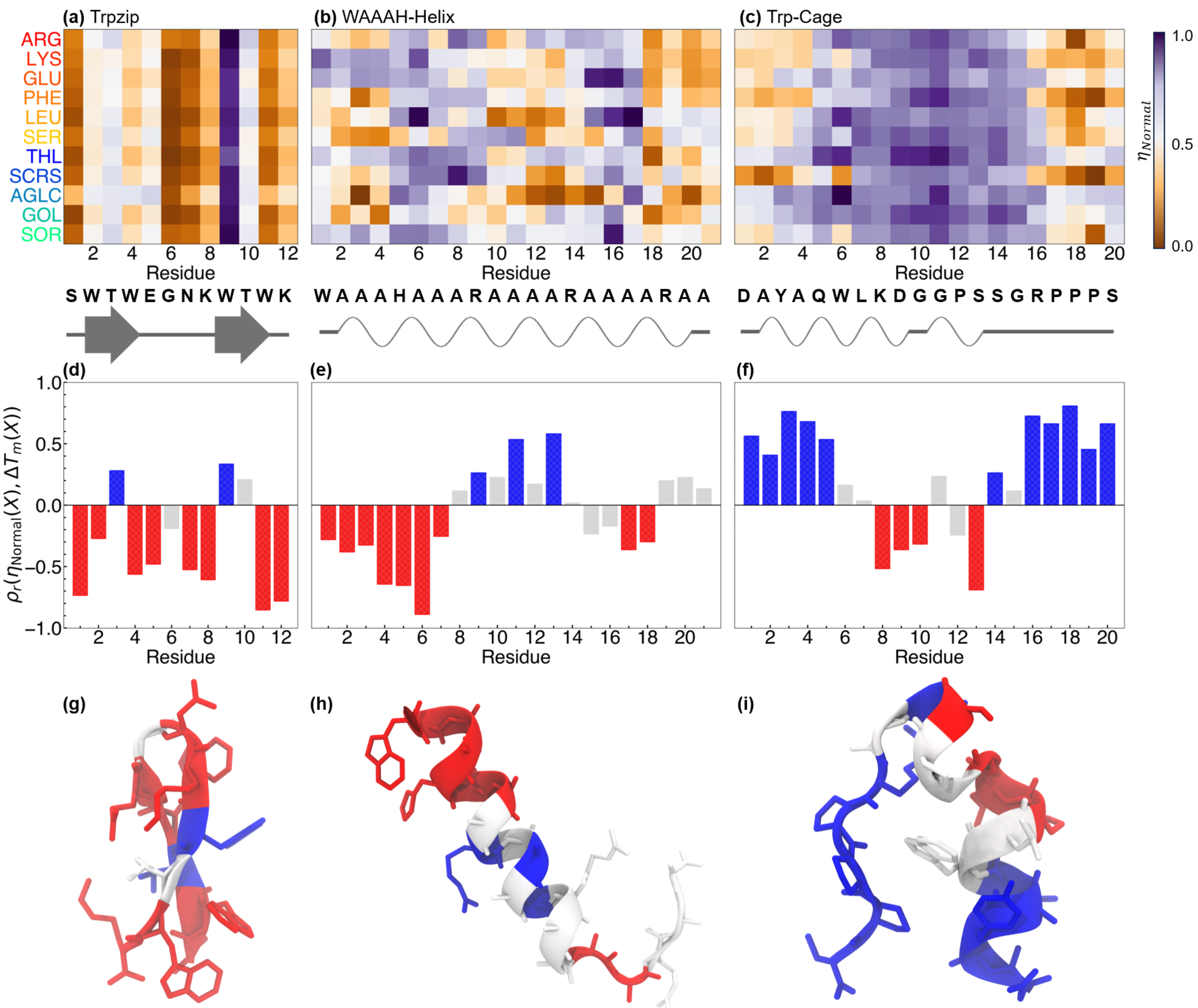


Figure S14: Per-residue normal similarity scores, $\eta_{Normal}$. (a-c) Heatmaps are shown for proteins in different excipient solutions, (a) Trpzip, (b) WAAAH-Helix, (c) Trp-Cage. Below each plot, the amino acid sequence is provided according to canonical one-letter codes. Secondary structure assignment was determined by DSSP[S27] and plotted using Biotite,[S28] where $\beta$-sheets are drawn as arrows and $\alpha$-helices are springs. $\boldsymbol{\eta}$ is normalized from 0 (orange) to 1 (purple) in (a-c). (d-f) Spearman correlation coefficients, $\rho$, of $\eta$ with $\Delta T_m$ for (d) Trpzip, (e) WAAAH-Helix, and (f) Trp-Cage. Colors represent negative correlation (red) and positive correlation (blue). Residues where $|\rho| < 0.25$ are colored gray. (g-i) $\rho$ values are projected onto representative configurations of (g) Trpzip, (h) WAAAH-Helix, and (i) Trp-Cage.

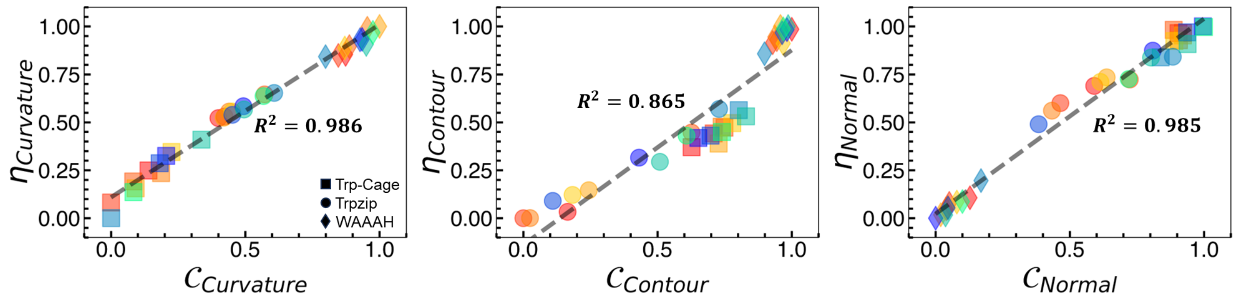


Figure S15: Correlation and $R^2$ values for local and global shape complementarity scores. (a) Curvature, (b) contour, and (c) normal. Data correspond to the temperature replica closest to the melting temperature of each protein in water (e.g., REST3 index 2). Different proteins are indicated by different shapes (Trp-Cage, square; Trpzip, circle; WAAAH, triangle), while excipients are indicated by the coloring scheme provided in Fig. 2.

#### 2.4 Normalization and Error Estimates

Calculations of $\mathcal{C}_{Curvature}$, $\mathcal{C}_{Contour}$, and $\mathcal{C}_{Normal}$ are theoretically bounded from 0–1 by construction, though in practice we observed their actual distributions to be narrow. To remove differences in scale and spread across metrics, each component $\mathcal{C}_i$ is normalized across all excipient/protein combinations at a given effective temperature using min-max normalization. This additionally improves the stability and interpretability of multivariate regression (see Section 2.5), which optimizes the relationship between $\Omega$ and $\Delta T_m$. Composite scores, $\Omega$, are additionally normalized using min-max normalization to aid in visualization and interpretability of the data. The same normalization procedure is applied to local shape complementarity scores, $\eta_i$.

For each global shape complementarity metric, a time-dependent quantity is computed as $\mathcal{C}_i(t)$. Mean values are reported as the time-averaged values, $\langle C_i(t)\rangle = \frac{1}{T}\sum_{t=0}^{T}\mathcal{C}_i(t)$, where $t$ is a simulation frame at time $t$ and $T$ is the total number of frames. Errors are reported as standard error from the mean value, $SEM = \sigma_i/\sqrt{T}$. For local shape complementarity metrics, a pooled estimator is calculated over all observations across frames and vertices corresponding to residue $j$:

$$\langle \eta_{i,j}\rangle = \frac{1}{N_j}\sum_{k=1}^{N_j}\eta_{i,j}^{(k)} \tag{S7}$$

where $N_j$ is the total number of observations for residue $j$. In this case, the standard error estimate becomes $SEM = \sigma_i/\sqrt{N_j}$.

#### 2.5 Linear Regression and Sigmoidal Fine-Tuning

The composite scoring function $\Omega$ depends on prior calculation of weights, $\omega_i$. These are computed using a multivariate regression, performed to optimize the relationship between shape complementarity, $\Omega$, and the change in melting temperature, $\Delta T_m$:

$$\Omega = \sum_i \omega_i \mathcal{C}_i \tag{S8}$$

$$X = [\mathcal{C}_1 \ \mathcal{C}_2 \ \mathcal{C}_3 \ \mathbf{1}] \tag{S9}$$

$$y = \Delta T_m \tag{S10}$$

$$\beta = \begin{bmatrix} \omega_1 \\ \omega_2 \\ \omega_3 \\ b \end{bmatrix} \tag{S11}$$

$$\hat{\beta} = (X^\top X)^{-1} X^\top y \tag{S12}$$

$$\hat{y} = X\hat{\beta} = \Delta\hat{T}_m = \omega_1\mathcal{C}_1 + \omega_2\mathcal{C}_2 + \omega_3\mathcal{C}_3 + \beta_0 \tag{S13}$$

The coefficient of determination is determined according to:

$$R^2 = 1 - \frac{\sum_i (y_i - \hat{y}_i)^2}{\sum_i (y_i - \bar{y})^2} \tag{S14}$$

And weights are rescaled according to:

$$\hat{\omega_i} = \frac{\omega_i}{\sum_i \omega_i} \tag{S15}$$

to match the normalization of $\Omega$. Following linear regression, the weights are fixed to compute $\Omega$. $\Delta T_m$ and $\Omega$ are then used to fit a sigmoidal function as a fine-tuning measure upon visual inspection of the relationship among the data. The sigmoid function is defined as:

$$\Omega(\Delta\hat{T}_m) = \frac{1}{1 + e^{-a\Delta\hat{T_m}+b}} \quad \text{(S16)}$$

where $a$ and $b$ are parameters that are fit to the data. A summary of optimized weights and $R^2$ values obtained from multivariate regression and lines of best fit and sigmoidal fine-tuning for miniprotein systems are provided in Table S6. Data for larger, globular protein are provided in Table S7.

Table S6: Optimized weights and regression metrics for each REST3 replicate simulation. Effective temperatures corresponding to REST3 indices (i) are provided for Trp-Cage and Trpzip ($T_{Eff}^{TC/TZ}$) and WAAAH-Helix ($T_{Eff}^{WH}$).

| i | $T_{Eff}^{TC/TZ}$ [K] | $T_{Eff}^{WH}$ [K] | $\omega_1$ | $\omega_2$ | $\omega_3$ | a | b | $R^2_{Linear}$ | $R^2_{Sigmoid}$ |
|---|---|---|---|---|---|---|---|---|---|
| 0 | 300.000 | 260.000 | 0.7526 | 0.3791 | -0.1317 | 0.071 | -0.096 | 0.7265 | 0.8410 |
| 1 | 313.944 | 273.425 | 0.6516 | 0.2845 | 0.0639 | 0.087 | -0.077 | 0.7627 | 0.9137 |
| 2 | 328.708 | 287.727 | 0.6594 | 0.1683 | 0.1723 | 0.077 | -0.187 | 0.7954 | 0.9254 |
| 3 | 344.351 | 302.977 | 0.7162 | 0.0020 | 0.2818 | 0.045 | -0.188 | 0.8102 | 0.8744 |
| 4 | 360.934 | 319.251 | 0.7546 | -0.0533 | 0.2987 | 0.052 | -0.899 | 0.7596 | 0.8809 |
| 5 | 378.529 | 336.632 | 0.7239 | 0.1027 | 0.1734 | 0.055 | -1.177 | 0.7434 | 0.9054 |
| 6 | 397.208 | 355.213 | 0.6249 | 0.5252 | -0.1501 | 0.056 | -1.281 | 0.7184 | 0.8835 |
| 7 | 417.055 | 375.093 | 0.5678 | 0.0696 | 0.3626 | 0.050 | -1.286 | 0.7405 | 0.8926 |
| 8 | 438.156 | 396.383 | 0.4984 | 0.2423 | 0.2593 | 0.063 | -1.053 | 0.6694 | 0.8363 |
| 9 | 460.609 | 419.206 | 0.4630 | 0.5932 | -0.0562 | 0.103 | -1.459 | 0.6493 | 0.8470 |
| 10 | 484.519 | 443.696 | -0.6179 | 1.9254 | -2.3075 | 0.114 | -1.016 | 0.6345 | 0.8293 |
| 11 | 510.000 | 470.000 | 0.3331 | 1.4709 | -0.8039 | 0.128 | -1.229 | 0.6329 | 0.8370 |

#### 2.6 Extension to Globular Proteins

An additional 12 replicate simulations were performed for globular proteins hen egg white lysozyme (HEWL) and thaumatin (TMT). For miniproteins, we observed the strongest correlation between $\Delta T_m$ and $\Omega$ near the melting temperature of each miniprotein in water. To approximate the conformational ensemble of HEWL and TMT near their melting temperatures, we generated plausible folding intermediates by performing short replica exchange umbrella sampling (REUS)[S29] simulations along the fraction of native contacts (Q) reaction coordinate. The final configuration from each of 12 evenly spaced windows in the range

$0 \leq Q \leq 1$ were used as starting configurations for protein-excipient equilibrium simulations. 0.1 M excipients (ARG, LYS, GLU, THL, SCRS, and SOR) were added to each initial protein configuration. Equilibration and production runs were then carried out as described previously. Proteins were then simulated near their experimental melting temperatures in water (337 K for TMT; 343 K for HEWL). RMSD values and representative snapshots showing the conformational diversity of these simulations are shown in Figs. S16 and S17. Radius of gyration $R_g$ values are provided in Figs. S18 and S19. The correlation between $\Omega$ and $\Delta T_m$ following multivariate regression when considering only simulations initialized from $Q \approx 1$ or $Q \approx 0$ are shown in Fig. S20.

Table S7: Optimized weights and regression metrics for simulations of hen egg white lyzoyme (HEWL) and thaumatin (TMT).

| Condition | $\omega_1$ | $\omega_2$ | $\omega_3$ | a | b | $R^2_{Linear}$ | $R^2_{Sigmoid}$ |
|---|---|---|---|---|---|---|---|
| Equivalent Weights | 0.333 | 0.333 | 0.333 | 0.129 | 0.261 | 0.008 | 0.008 |
| Miniprotein Weights | 0.659 | 0.168 | 0.077 | -0.499 | -0.123 | 0.112 | 0.108 |
| Optimized Weights, All Excipients | -1.405 | -2.492 | 2.897 | 2.983 | 1.182 | 0.838 | 0.910 |
| Optimized Weights, Amino Acid Excipients | -0.324 | -0.774 | 0.098 | 2.587 | 1.076 | 0.822 | 0.843 |
| Optimized Weights, Sugar Excipients | -0.332 | 0.224 | 1.108 | 3.838 | 0.648 | 0.991 | 0.991 |

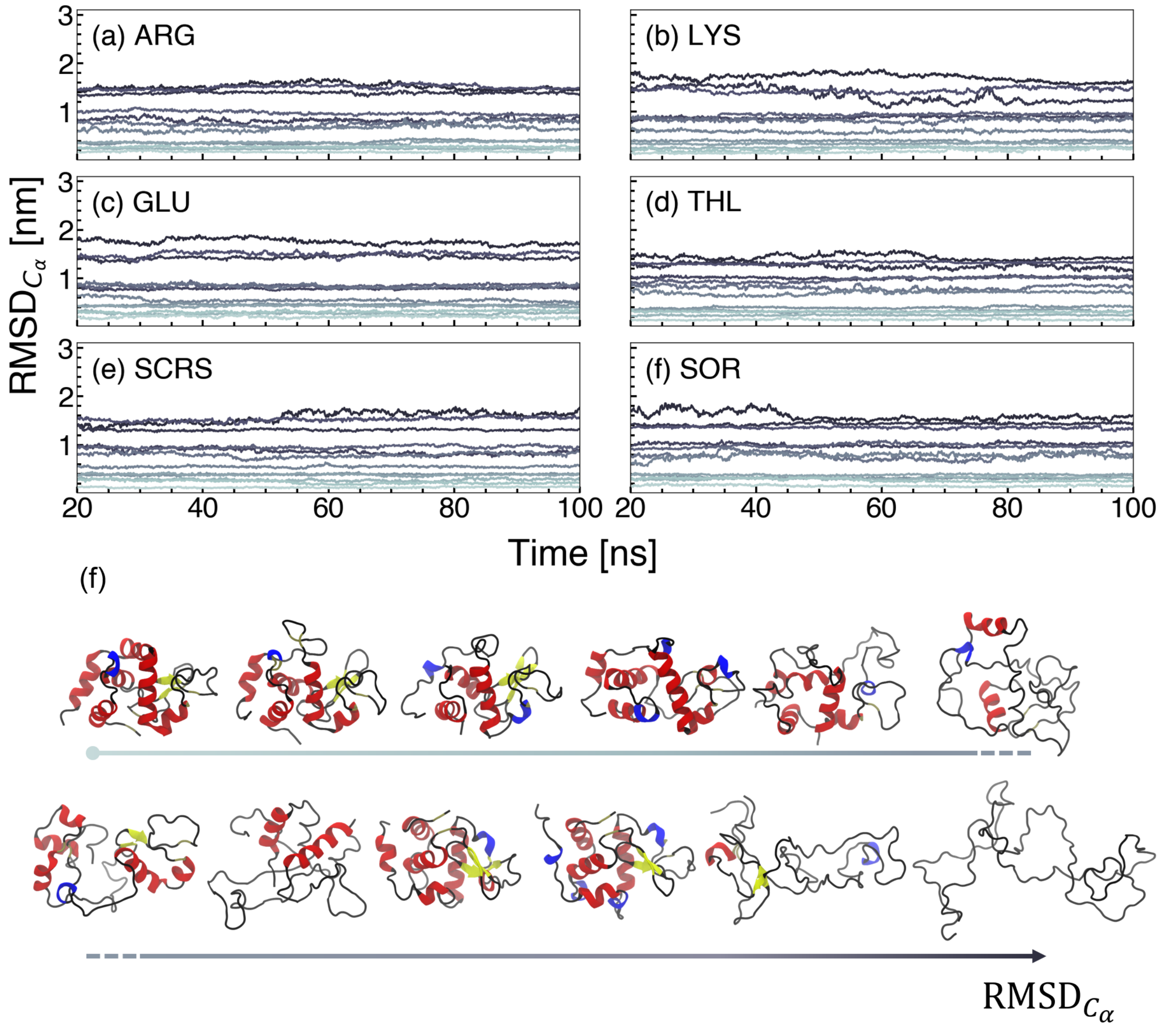


Figure S16: RMSD$_{C_\alpha}$ versus time in HEWL/excipient simulations. Each protein/excipient pair is run for 100 ns in 12 independent simulations, where each simulation has a different initial configuration extracted from REUS simulations along Q. (a) ARG, (b) LYS, (c) GLU, (d) THL, (e) SCRS, (f) SOR. (f) Representative snapshots from simulations with increasing RMSD$_{C_\alpha}$ values.

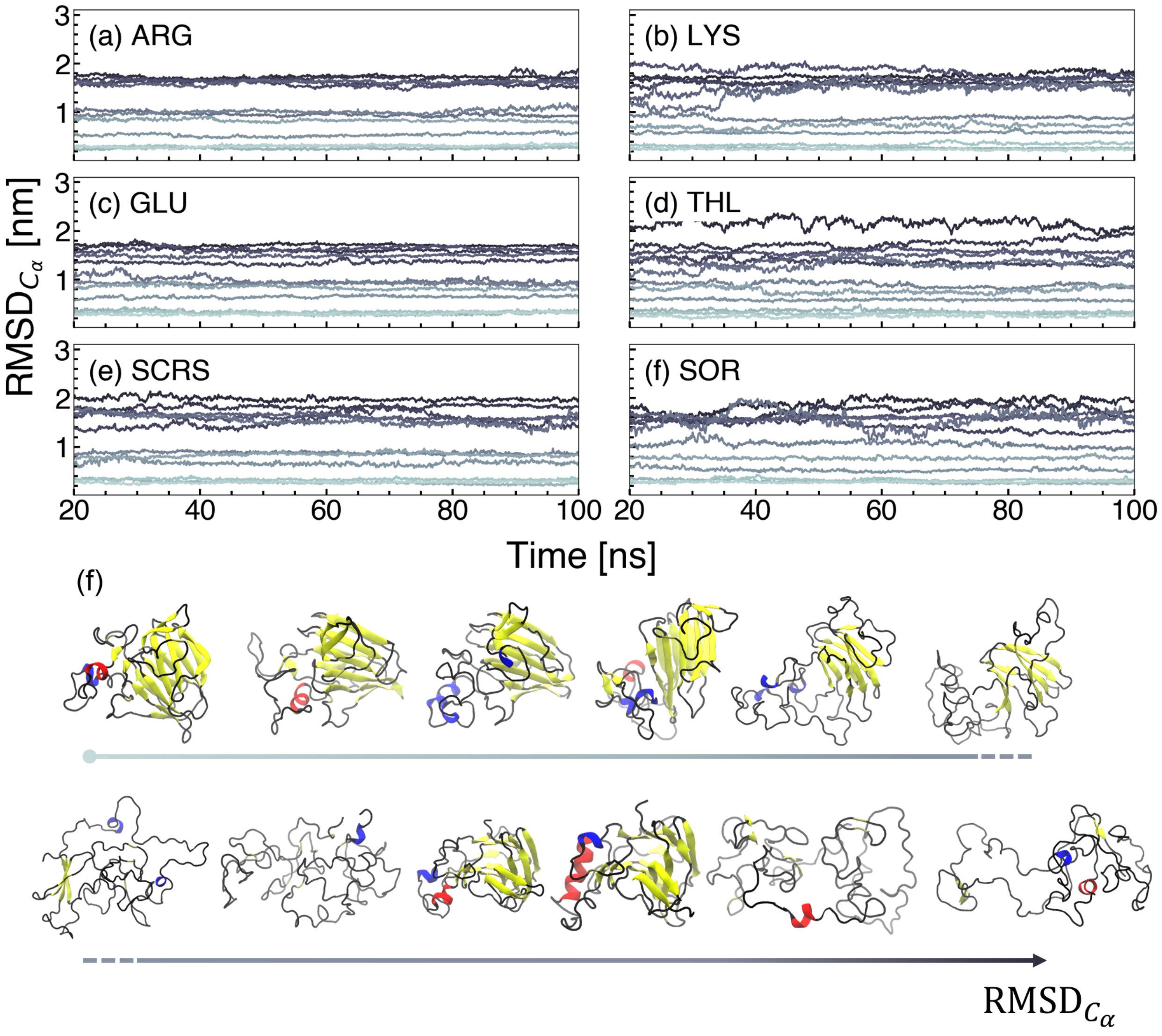


Figure S17: RMSD$_{C_\alpha}$ versus time in TMT/excipient simulations. Each protein/excipient pair is run for 100 ns in 12 independent simulations, where each simulation has a different initial configuration extracted from REUS simulations along Q. (a) ARG, (b) LYS, (c) GLU, (d) THL, (e) SCRS, (f) SOR. (f) Representative snapshots from simulations with increasing RMSD$_{C_\alpha}$ values.

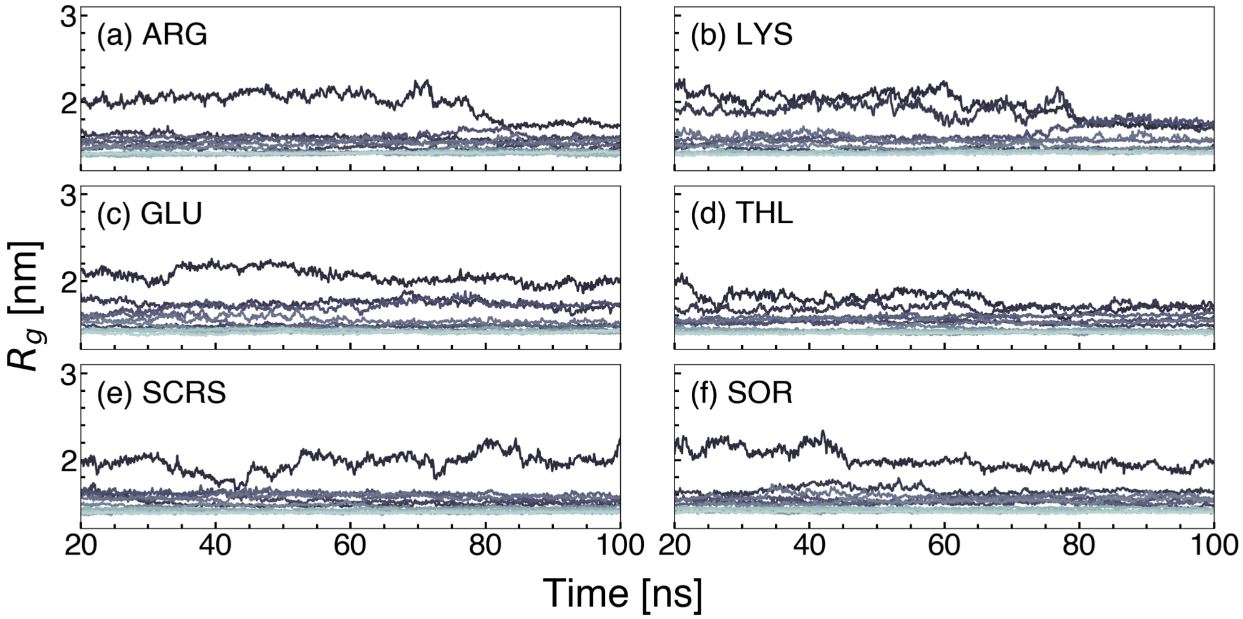


Figure S18: $R_g$ versus time in HEWL/excipient simulations. Each protein/excipient pair is run for 100 ns in 12 independent simulations, where each simulation has a different initial configuration extracted from REUS simulations along Q. (a) ARG, (b) LYS, (c) GLU, (d) THL, (e) SCRS, (f) SOR.

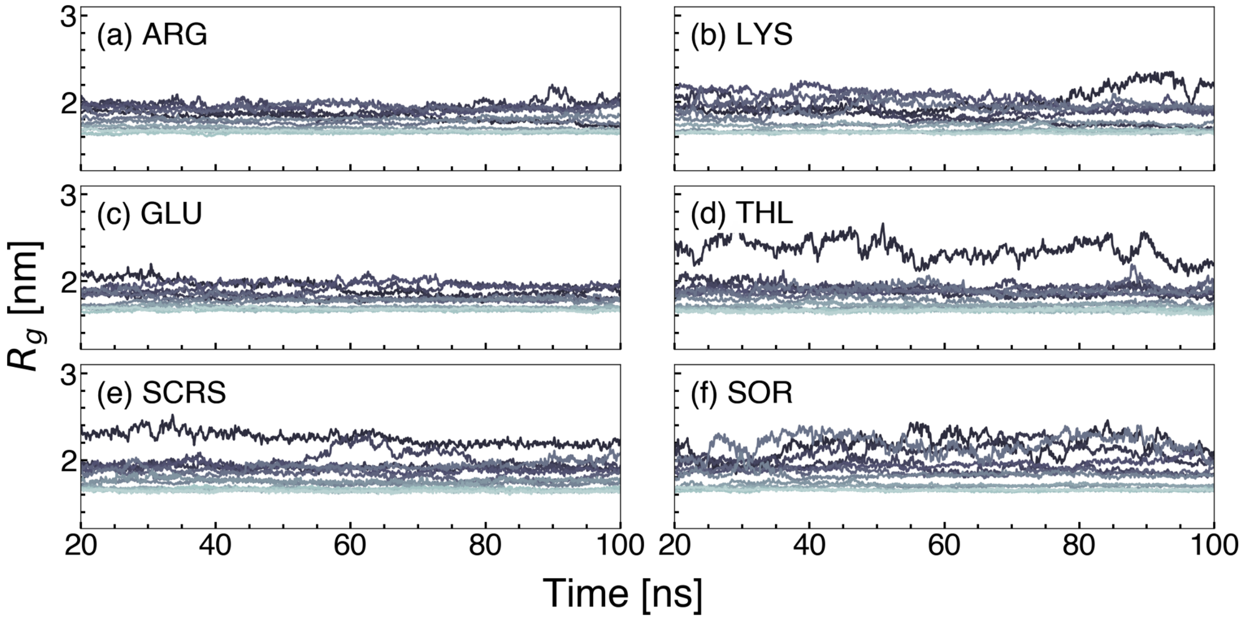


Figure S19: $R_g$ versus time in TMT/excipient simulations. Each protein/excipient pair is run for 100 ns in 12 independent simulations, where each simulation has a different initial configuration extracted from REUS simulations along Q. (a) ARG, (b) LYS, (c) GLU, (d) THL, (e) SCRS, (f) SOR.

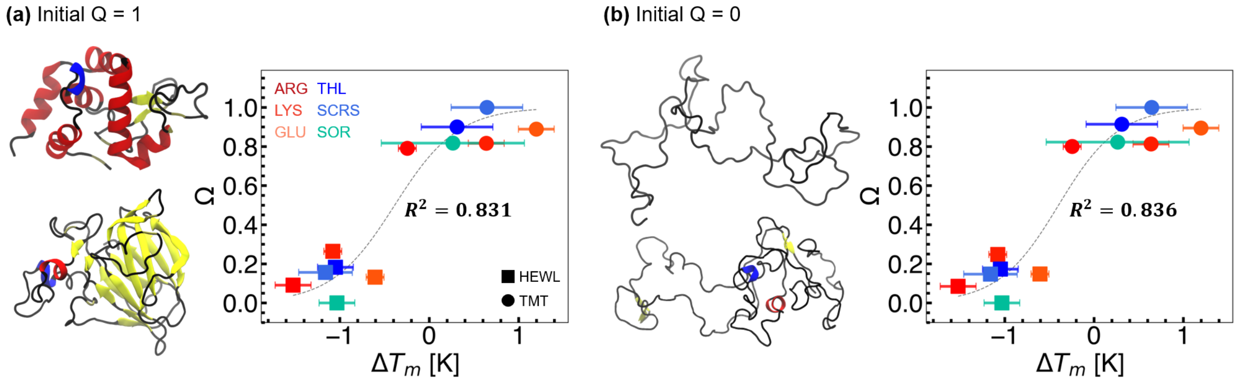


Figure S20: Relationship between optimized $\Omega$ values and $\Delta T_m$ when considering only simulations initialized with configurations where (a) $Q \approx 1$ or (b) $Q \approx 0$.

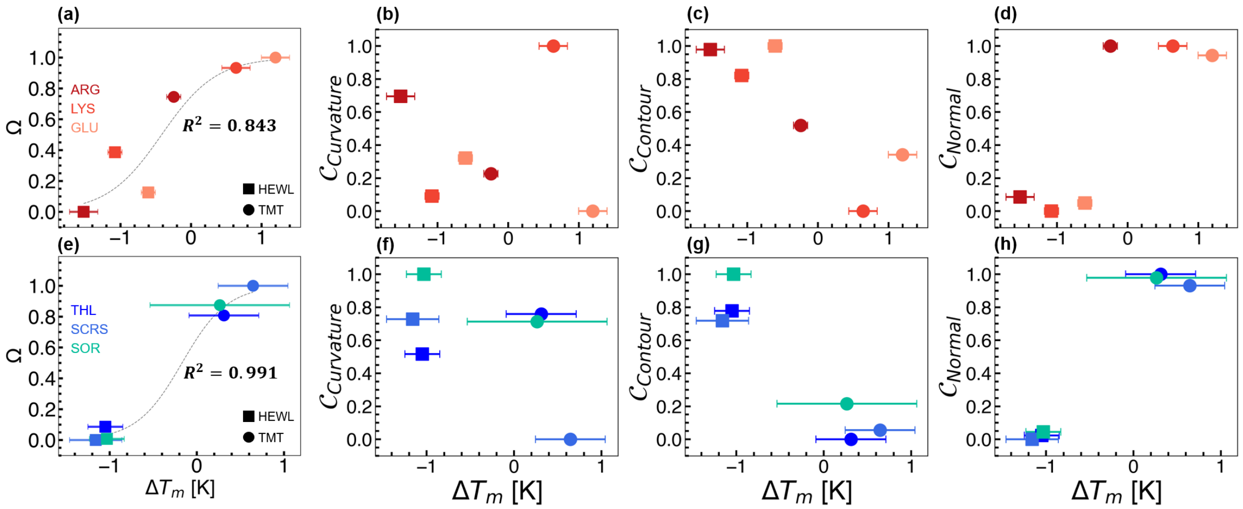


Figure S21: Relationships between protein-solvent shape complementarity and change in the temperature stability of larger, globular proteins. (a-d) Scatter plots showing shape complementarity versus change in melting temperature for amino acid excipients, (a) $\Omega$, (b) $\mathcal{C}_{Curvature}$, (c) $\mathcal{C}_{Contour}$, (d) $\mathcal{C}_{Normal}$. (e-h) Scatter plots showing shape complementarity versus change in melting temperature for sugar excipients, (e) $\Omega$, (f) $\mathcal{C}_{Curvature}$, (g) $\mathcal{C}_{Contour}$, (h) $\mathcal{C}_{Normal}$. Different proteins are indicated by different shapes (HEWL, square; TMT, circle), while excipients are indicated by different colors as in Fig. 2. Dashed lines represent the sigmoidal fit following fine-tuning.

#### 2.7 Pseudocodes

**Algorithm 1** High-Level Overview

```
for frame in trajectory do
    1. Extract protein heavy atoms
    2. Extract solvent heavy atoms in shell [rThresh2, rThresh1]
    3. Build surfaces:
         protein_surface ← mesh(protein_coords)
         solvent_surface ← mesh(solvent_coords)
    4. Inflate/deflate surfaces using van der Waals radii:
         protein_surface ← push outward along normals
         solvent_surface ← push inward along normals
    5. Compute global shape metrics:
         - contour similarity
         - curvature similarity
         - normal similarity
    6. Compute residue-level contributions:
         - local curvature
         - local distance
         - local normal alignment
    7. Store results
```

**Algorithm 2** Shape Similarity Metrics

```
procedure Contour Similarity
    for vertex p on protein_surface do
        d(p) ← distance to nearest point on solvent_surface
    mean_d = mean(d)
    stdv_d = std(d)
    distance_variation = stdv_d / mean_d
    contour_similarity = exp(-distance_variation)
procedure Curvature Similarity
    curvature_protein ← Gaussian curvature at all protein_surface vertices
    curvature_solvent ← Gaussian curvature at all solvent_surface vertices
    emd = WassersteinDistance(curvature_protein, curvature_solvent)
    curvature_similarity = exp(-emd)
procedure Normal Similarity
    Sample N points on protein_surface
    for each sampled point p do
        q ← closest point on solvent_surface
        n_p ← normal at p
        n_q ← normal at q
        cos_theta(p) = dot(n_p, n_q) / (|n_p| * |n_q|)
    mean_alignment = mean(cos_theta over all samples)
    normal_similarity = (1 + mean_alignment) / 2
```

### 3 Experimental Temperature Stability Assays

#### 3.1 Materials

Hydrochloric acid (HCl, ACS grade), sodium hydroxide (NaOH, ACS grade), sodium phosphate monobasic anhydrous ($NaH_2PO_4$, ≥95%), and sucrose (>95%) were purchased from Fisher Scientific. D-(+)-trehalose (trehalose, 99%), D-sorbitol (sorbitol, 100%), and D-(+)-glucose monohydrate (glucose, 99%) were purchased from Alfa Aesar Thermo Fisher Scientific Chemicals. L-lysine hydrochloride (Lys, Ultrapure) was purchased from Thermo Fisher Scientific. L-glutamic acid monosodium salt (Glu, cell culture reagent) was purchased from MP Biomedicals. Glycerol (99+%) was purchased from Acros Organics. L-arginine hydrochloride (Arg, ≥ 99.0%) and Thaumatin from *Thaumatococcus daniellii* (TMT) were purchased from Sigma-Aldrich. Hen egg white lysozyme (HEWL, ≥95%) was purchased from Hampton Research. Sypro Orange was purchased as a 50x solution as a Protein Thermal Shift™ Dye Kit (PTSD Kit, Applied Biosystems). Deionized (DI) water was obtained from a Milli-Q water purification system (resistivity of 18.2 MΩ cm, Millipore).

#### 3.2 Stock Solution Preparation

Stock solutions of 1 M NaOH and 1 M HCl were prepared gravimetrically in DI water. Phosphate buffer (PB, pH $\sim$ 7.2) containing 1.54 mM $NaH_2PO_4$ and 2.71 mM $NaH_2PO_4$ were prepared gravimetrically in DI water. Excipient stock solutions at 0.1 M concentration were prepared with PB. 50x Sypro Orange stock solutions were prepared with the dye and the buffer from the PTSD Kit in a volumetric ratio of 1 to 49.

#### 3.3 Sample Preparation

Samples were prepared by pipetting the relevant buffer, excipient, protein, and Sypro Orange, in this order, into a microcentrifuge tube (Fisher Scientific) for manual sample preparation or into a 96-well plate (Falcon) via liquid handling robot (Biomek NX$^P$ Beckman Coulter). The sample was mixed via vortexing after the addition of protein and again after adding Sypro Orange. The ratio of protein to dye was optimized for each protein to ensure repeatable non-negligible peaks in first derivative curves (details are in the characterization section). The volume ratio of 50x Sypro Orange stock solution to protein stock solution to the entire test sample volume was fixed as 8:7:65 for HEWL and 8:7:65 for TMT in PB. Samples prepared manually were at a total volume of 80 $\mu$L while samples prepared using the liquid handling robot had a total volume of 160 $\mu$L.

#### 3.4 Thermal Shift Characterization

Differential scanning fluorimetry (DSF) using the Sypro Orange dye was used to measure protein stability as described by a hydrophobic exposure temperature (HET),[S30,S31] which has been shown to have a strong correlation with the melting temperature of proteins. DSF experiments were performed by pipetting three 25 $\mu$L replicate aliquots into a 96-well PCR plate (Thermo Scientific). The PCR plate was gently centrifuged to eliminate bubbles by pulsing the centrifugation in a Thermo Scientific SORVALL ST 16R for 5 seconds before being loaded into a CFX Connect RT-PCR (Bio-Rad) system to perform DSF. Fluorescence intensity was collected over the temperature range of 10 °C to 95 °C in 1 °C increments in FRET mode (i.e., excitation wavelengths of 450 nm to 490 nm and detection wavelengths of 560 nm to 580 nm), with each increment stabilizing for 1 minute before the measurements were taken.

The HET is typically defined as the temperature at the DSF melting curve inflection point, determined by the negative first derivative, as described in Eq. S17. The HET was determined by identifying the minimum value of the first derivative and then performing a quadratic fit using two points on either side of the numerical minimum. The reported value for the HET is the average of three or more observations, and error bars express the standard deviation.

Table S8: Hydrophobic exposure temperature (HET) data for proteins with 0.1 M excipients. Data are shown in °C. Errors are reported as the standard deviation from three replicate samples.

| **Solution** | **Sample 1** | **Sample 2** | **Sample 3** | **Average** | **Error** |
|---|---|---|---|---|---|
| HEWL / Water | 70.0 | 69.8 | 69.8 | 69.9 | 0.1 |
| HEWL / ARG | 68.3 | 68.3 | 68.4 | 69.0 | 0.0 |
| HEWL / LYS | 68.8 | 68.8 | 68.8 | 68.8 | 0.0 |
| HEWL / GLU | 69.2 | 69.3 | 69.2 | 69.2 | 0.1 |
| HEWL / THL | 68.9 | 68.8 | 68.7 | 68.8 | 0.1 |
| HEWL / SCRS | 68.5 | 68.7 | 68.8 | 68.7 | 0.2 |
| HEWL / SOR | 68.8 | 68.9 | 68.9 | 68.8 | 0.1 |
| TMT / Water | 63.7 | 63.8 | 63.8 | 63.8 | 0.1 |
| TMT / ARG | 63.5 | 63.5 | 63.5 | 63.5 | 0.0 |
| TMT / LYS | 64.5 | 64.4 | 64.4 | 64.4 | 0.1 |
| TMT / GLU | 64.9 | 65.1 | 64.9 | 65.0 | 0.1 |
| TMT / THL | 64.8 | 63.9 | 63.8 | 64.1 | 0.4 |
| TMT / SCRS | 64.2 | 64.3 | 64.7 | 64.4 | 0.3 |
| TMT / SOR | 64.3 | 64.6 | 63.2 | 64.0 | 0.7 |

$$HET = -\frac{\partial(Fluorescence\ Response)}{\partial T} \tag{S17}$$

To quantitatively determine the performance of excipients on stabilizing proteins in fixed buffer solutions, we estimate $\Delta T_m$ experimentally according to Eq. S18.

$$\Delta T_m \approx \Delta HET = HET_{Excipient} - HET_{Buffer} \tag{S18}$$

(S11) Lorentz, H. A. Ueber die Anwendung des Satzes vom virial in der Kinetischen Theorie der Gase. *Annalen der Physik* **1881**, *248*, 127–136.

(S12) Berthelot, D. Sur le mélange des gaz. *Compt. Rendus* **1898**, *126*.

(S13) Brooks, B. R.; Brooks, C. L.; Mackerell, A. D.; Nilsson, L.; Petrella, R. J.; Roux, B.; Won, Y.; Archontis, G.; Bartels, C.; Boresch, S. et al. CHARMM: The biomolecular simulation program. *J. Comput. Chem.* **2009**, *30*, 1545–1614.

(S14) Huang, J.; MacKerell, A. D. CHARMM36 all-atom additive protein force field: Validation based on comparison to NMR data. *J. Comput. Chem.* **2013**, *34*, 2135–2145.

(S15) Cloutier, T.; Sudrik, C.; Sathish, H. A.; Trout, B. L. Kirkwood–Buff-derived alcohol parameters for aqueous carbohydrates and their application to preferential interaction coefficient calculations of proteins. *The Journal of Physical Chemistry B* **2018**, *122*, 9350–9360.

(S16) Muralikrishnan, P.; Zajac, J. W. P.; Heldt, C. L.; Perry, S. L.; Sarupria, S. Thermodynamic Basis of Sugar-Dependent Polymer Stabilization: Informing Biologic Formulation Design. *J. Phys. Chem. B* **2025**, *129*, 13018–13031.

(S17) Jorgensen, W. L.; Chandrasekhar, J.; Madura, J. D.; Impey, R. W.; Klein, M. L. Comparison of simple potential functions for simulating liquid water. *The Journal of Chemical Physics* **1983**, *79*, 926–935.

(S18) Cochran, A. G.; Skelton, N. J.; Starovasnik, M. A. Tryptophan zippers: Stable, monomeric -hairpins. *Proc. Natl. Acad. Sci. U.S.A.* **2001**, *98*, 5578–5583.

(S19) Barua, B.; Lin, J. C.; Williams, V. D.; Kummler, P.; Neidigh, J. W.; Andersen, N. H. The Trp-cage: optimizing the stability of a globular miniprotein. *Protein Engineering Design and Selection* **2008**, *21*, 171–185.

(S20) Talon, R.; Nauton, L.; Canet, J.-L.; Kahn, R.; Girard, E.; Gautier, A. Clicked europium dipicolinate complexes for protein X-ray structure determination. *Chem. Commun.* **2012**, *48*, 11886.

(S21) Hanwell, M. D.; Curtis, D. E.; Lonie, D. C.; Vandermeersch, T.; Zurek, E.; Hutchison, G. R. Avogadro: an advanced semantic chemical editor, visualization, and analysis platform. *J Cheminform* **2012**, *4*, 17.

(S22) Olsson, M. H. M.; Søndergaard, C. R.; Rostkowski, M.; Jensen, J. H. PROPKA3: Consistent Treatment of Internal and Surface Residues in Empirical p $K_{\mathrm{a}}$ Predictions. *J. Chem. Theory Comput.* **2011**, *7*, 525–537.

(S23) Zhang, Y.; Liu, X.; Chen, J. Re-Balancing Replica Exchange with Solute Tempering for Sampling Dynamic Protein Conformations. *J. Chem. Theory Comput.* **2023**, *19*, 1602–1614.

(S24) Liu, P.; Kim, B.; Friesner, R. A.; Berne, B. J. Replica exchange with solute tempering: A method for sampling biological systems in explicit water. *Proc. Natl. Acad. Sci. U.S.A.* **2005**, *102*, 13749–13754.

(S25) Wang, L.; Friesner, R. A.; Berne, B. J. Replica Exchange with Solute Scaling: A More Efficient Version of Replica Exchange with Solute Tempering (REST2). *J. Phys. Chem. B* **2011**, *115*, 9431–9438.

(S26) Barber, C. B.; Dobkin, D. P.; Huhdanpaa, H. The quickhull algorithm for convex hulls. *ACM Trans. Math. Softw.* **1996**, *22*, 469–483.

(S27) Frishman, D.; Argos, P. Knowledge-based protein secondary structure assignment. *Proteins* **1995**, *23*, 566–579.

(S28) Kunzmann, P.; Hamacher, K. Biotite: A Unifying Open Source Computational Biology Framework in Python. *BMC Bioinformatics* **2018**, *19*, 346.

(S29) Sugita, Y.; Kitao, A.; Okamoto, Y. Multidimensional replica-exchange method for free-energy calculations. *J. Chem. Phys.* **2000**, *113*, 6042–6051.

(S30) Seabrook, S. A.; Newman, J. High-Throughput Thermal Scanning for Protein Stability: Making a Good Technique More Robust. *ACS Comb. Sci.* **2013**, *15*, 387–392.

(S31) Shi, S.; Semple, A.; Cheung, J.; Shameem, M. DSF Method Optimization and Its Application in Predicting Protein Thermal Aggregation Kinetics. *Journal of Pharmaceutical Sciences* **2013**, *102*, 2471–2483.
